# Infection cycle and phylogeny of the Polinton-like virus Phaeocystis globosa virus virophage-14T

**DOI:** 10.1101/2022.07.28.501842

**Authors:** Sheila Roitman, Andrey Rozenberg, Tali Lavy, Corina P. D. Brussaard, Oded Kleifeld, Oded Béjà

## Abstract

Virophages are small dsDNA viruses dependent on a nucleocytoplasmic large-DNA virus infection of a cellular host for replication. Putative virophages infecting algal hosts are classified together with polinton-like viruses, transposable elements widely found in algal genomes, yet the lack of isolated strains raises questions about their existence as independent entities. In this work we isolated and characterized a virophage (PgVV-14T) co-infecting *Phaeocystis globosa* with the Phaeocystis globosa virus-14T (PgV-14T). PgVV-14T decreases the fitness of its PgV-14T viral host, yet it does not salvage the cellular host population. We found viral-like elements resembling PgVV-14T in *Phaeocystis* genomes, suggesting that these virophages are capable of integrating to the cellular host genome, bridging the gap between Polinton-like viruses and virophages. This system, with a giant virus, a virophage and endogenous viral elements preying on an algal host, presents an opportunity to gain a better understanding on the evolution of eukaryotes and their viruses.

## Main

Eukaryotic genomes function as a hub where viruses and selfish genetic elements (SGEs) convene. This interplay between hosts and parasites leads to co-evolution, horizontal gene transfer, and genome diversification. Viruses and SGEs are capable of jumping in and out of the host genome; yet SGEs usually lack the structural proteins that could grant them independence from the host^1^. A remarkable group of Eukaryotic SGEs are the Polintons (or Mavericks), self-replicating large transposons^2–4^. Polintons are 15-20 kbp long, typically flanked by terminal inverted repeats (TIRs), and encode for a protein-primed DNA polymerase and a retrovirus-like integrase. Although there are no examples of Polintons becoming independent Polintoviruses, it has been proposed that Polintons are the ancestors of dsDNA viruses^4–7^. Recently, new groups of Polinton-like viruses (PLVs), who resemble Polintons, yet lack the polinton-distinctive genes, have been described by metagenomics; some PLVs were integrated into algal genomes, while others appear to be separated^8, 9^. To date, only one PLV particle has been isolated, TsV-N1, a small dsDNA nuclear-replicating virus infecting *Tetraselmis striata*^10^. PLVs encode a core module of three genes: a major and a minor capsid proteins (MCP, mCP) and a packaging-ATPase, along a variable set of genes conserved among Polintons and virophages^8^. PLVs associated with algae are classified in two groups, one of putative free-living PLVs represented by TsV-N1, and another including free and integrated PLVs, represented by *Phaeocystis globosa* virus virophage^8^.

Virophages are small viral parasites that depend on a helper virus from the nucleocytoplasmic large DNA viruses (NCLDV) for reproduction. Virophages are dsDNA viruses, have a 17-30-kbp genome with a low %GC content (27-50%) and icosahedral capsids of 35-70 nm in diameter^11, 12^. Most virophages were found by meta-genomics and remain uncultured^13–21^, yet a few were isolated conjointly with their helper from the *Mimiviridae* family^22–26^. Except for the recently isolated Chlorella virus virophage SW01^26^, all virophages have been classified within the *Lavidaviridae* family that currently contains two genera: *Sputnikvirus* and *Mavirus*^12^. Interestingly, Mavirus has homologous genes to Polintons^6, 23^ and integrated Mavirus-like elements were detected in *Cafeteria burkhardae* strains, indicating that maviruses lead a dual life-style, as free-living entities and integrated in the eukaryotic genome^16^.

In 2013, the genome of a virus infecting *Phaeocystis globosa,* PgV-16T was sequenced which led to an unexpected discovery of a co-occurring virophage genome in the assembly^27^. *Phaeocystis globosa* is a ubiquitous haptophyte capable of creating toxic blooms that can be terminated by viral infections^28, 29^. The genome of the PgV virophage (PgVV) was found to be ∼20 kbp, flanked by TIRs and low %GC content (36%)^27^. No small viral particles were observed along PgV-16T in viral lysates, and it was suggested that the virophage was packed as a linear plasmid within the PgV capsid or integrated in the viral genome^27^. Interestingly, three of PgVV’s encoded proteins had distant homologs in Mavirus, while the MCP, mCP and ATPase resembled those of algal PLVs^7, 8^. In this work, we report the isolation and characterization of Phaeocystis globosa virus virophage-14T, a PLV with a virophage life-style. We characterize the dynamics of PgV and PgVV infection of *Phaeocystis globosa,* and analyze virophages and Polinton-like genomes to create a framework to classify PLV virophages within the virosphere.

## Results and discussion

### PgV-14T and PgVV-14T sequencing

Within the framework of analyzing infection dynamics of *P. globosa* we sequenced the PgV-14T lysate. PgV-14T appeared almost identical to PgV-16T^27^, yet we found 16 single nucleotide polymorphisms, 6 indels and one small inversion (Supplementary Files 1, 3). The assembly also contained a separate scaffold corresponding to a genome closely related to the PgV-16T-associated PgVV^27^, that we designated PgVV-14T.

The PgVV-14T genome is syntenic with and highly similar to PgVV-16T, with minimal differences (for details see Supplementary Files 1, 3 and Extended Data Fig. 1). A deeper analysis of the genome revealed the presence of two previously omitted short ORFs, *pgvv01b* and *pgvv15b*. Contrary to the previous report^27^, we did not find any indication that PgVV-14T is integrated in PgV’s genome: out of 539,034 trimmed Illumina reads that could be mapped to the two reference genomes, no fragment was found to map to PgV and PgVV simultaneously.

**Figure 1.**
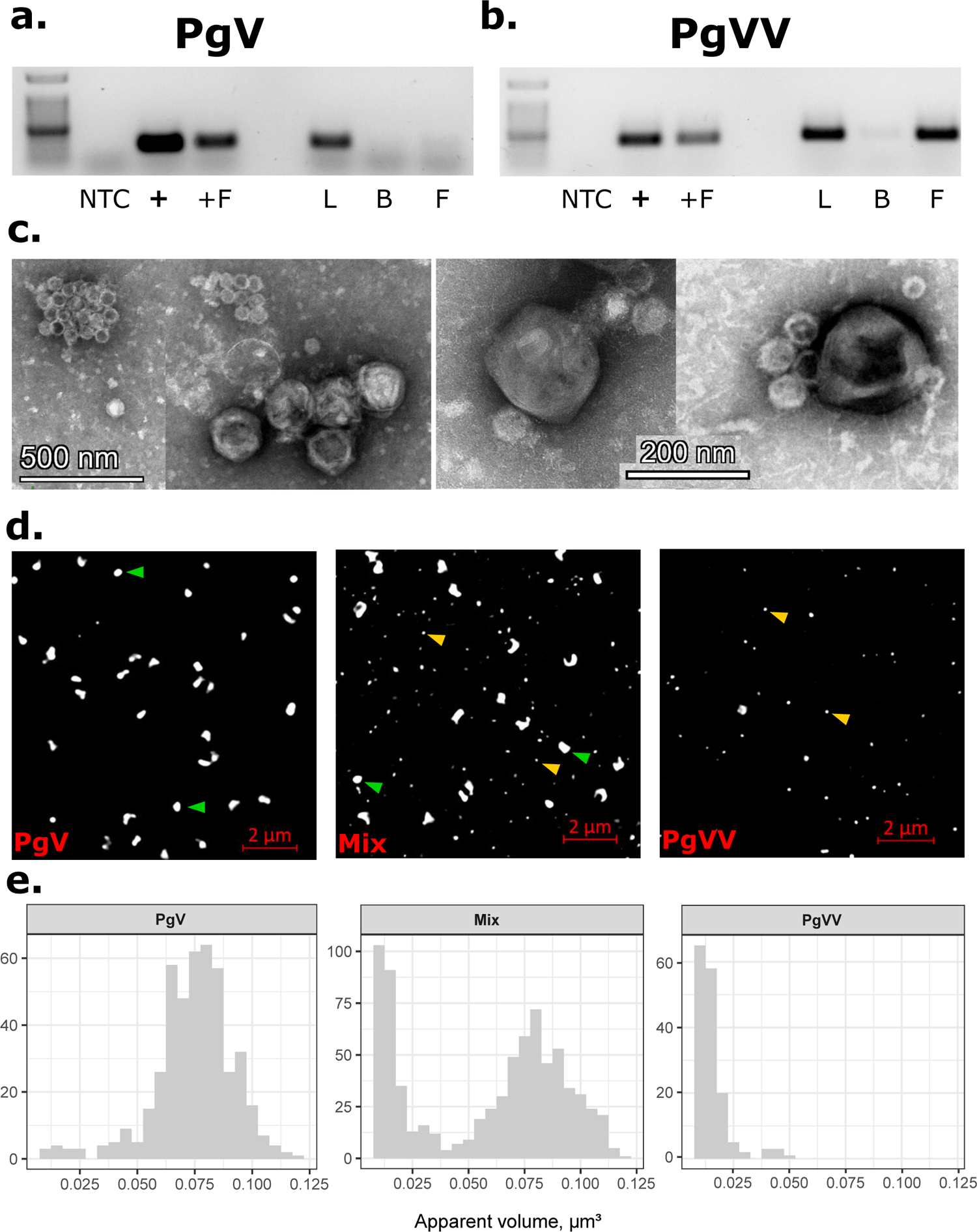
PgVV is a *bona-fide* virus. **a.** PCR assay with a PgV marker (MCP1, *pgv*157) and **b.** a PgVV marker (TVpol, *pgvv04*). NTC, No template control; +, Positive control (DNA); +F, 0.2 µm filtered DNA; L, lysate; B, boiled filtered lysate; F, 0.2 µm filtered lysate. L, B and F were subjected to DNAse treatment before the PCR. **c.** Transmission electron microscopy images of negatively stained PgV-only, PgV/PgVV mixed and PgVV-only lysates. **d.** SYBR green stained PgV only, mixed PgV and PgVV, and PgVV only lysates under Elyra 7 eLS microscope. Green and yellow arrows denote particles of PgV and PgVV size, respectively. **e.** Quantification of dots in SYBR green stained samples by apparent volume. Y-axis denotes the number of points counted.

### PgVV is a *bona-fide* virus

To assess whether the virophage genome is packed within PgV particles or in particles of its own, we filtered a fraction of a mixed viral lysate through a 0.2 µm filter (twice). Half of the filtrate was boiled and then all fractions (Lysate, Boiled and Filtered) were treated with DNAse to get rid of non-encapsidated DNA. PgVV marker genes could be amplified by PCR from both Lysate and Filtered fractions, while PgV was only found in the Lysate (Fig. 1a,b). Additionally, we prepared filters for viral counts by SYBR staining from PgV-only, PgVV-only and mixed lysates where we observed two distinct size populations matching the lysate expected composition (Fig. 1d,e). In line with this, under transmission electron microscopy (TEM), negatively stained lysates showed two types of icosahedral viral particles of different sizes (Fig. 1c). The larger particles measured 160-215 nm in diameter (mean 188 nm, s.d. 16 nm, n = 24), while the smaller particles measured 50-80 nm (mean 66 nm, s.d. = 7 nm, n = 77) (Supplementary File 3). Taken together, along with the proteomic identification of most PgVV proteins in purified viral lysates (see below), we conclude that PgVV-14T is a virus with a ∼19.5 kbp genome packed in icosahedral capsids of 66 nm.

### Infection dynamics

To get insight into the interactions between PgV and PgVV, we prepared a PgV-only lysate, by using a dilution-to-extinction approach; and a PgVV-only lysate, by filtering a mixed lysate through 0.1 µm filters followed by concentration and purification. *P. globosa* cultures were infected with either a PgV-only, a PgVV-only lysate or their mix. Cultures infected with PgV (solely or in conjunction with PgVV) were completely lysed after 3 or 4 days (respectively), while the culture infected with PgVV-only followed a similar growth pattern to the control (Extended Data Fig. 2, Supplementary File 2). PgV increased in abundance in both infection experiments, while PgVV increased only in the mixed infection. These results suggest that PgVV is unable to independently complete an infection cycle in *P. globosa*, and depends on PgV for its reproduction, the first experimental evidence that it leads a virophage lifestyle.

**Figure 2.**
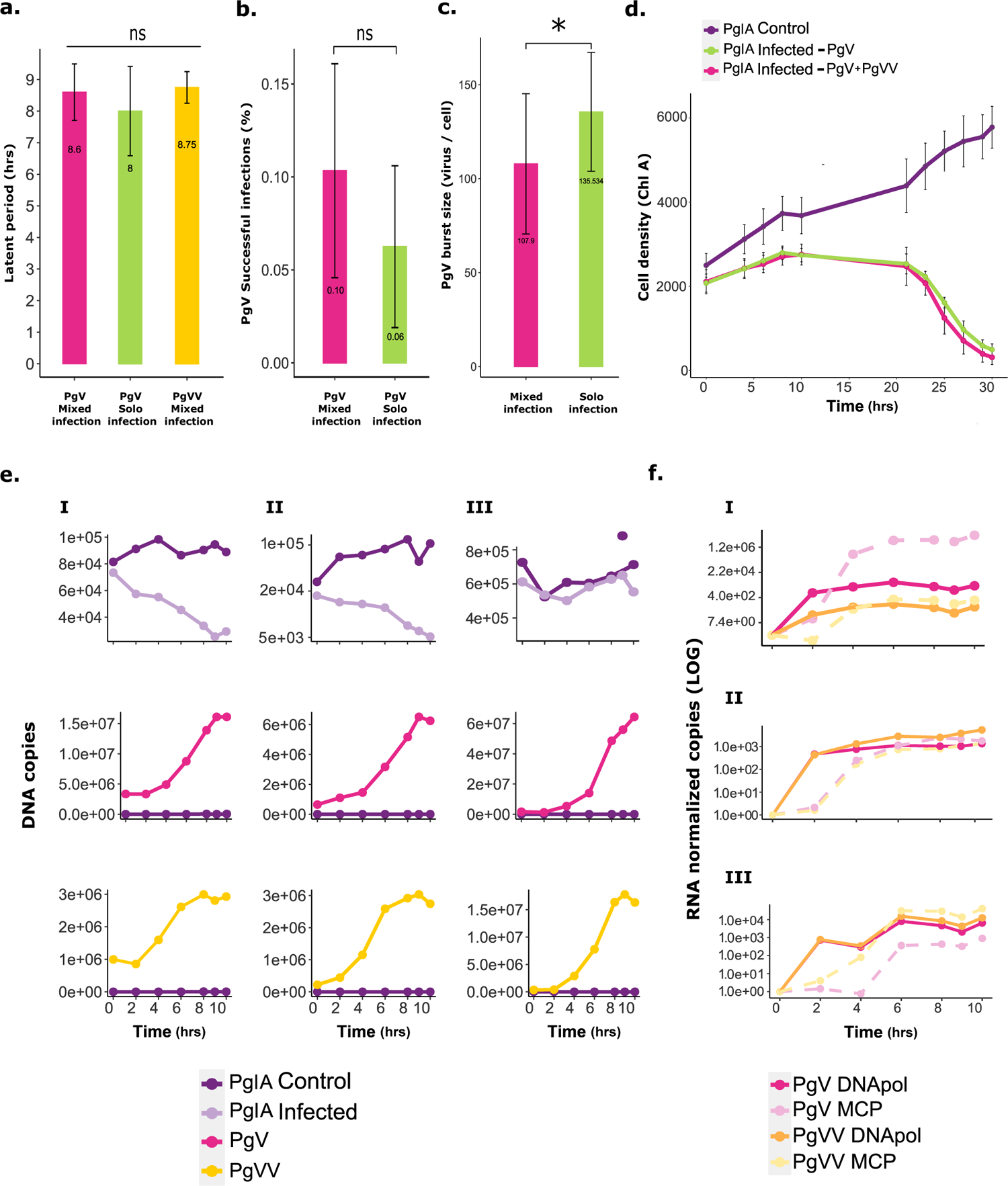
Infection dynamics of *P. globosa*, PgV and PgVV. **a.** Latent period of PgV-14T (Pink, in a mixed lysate with PgVV; Green, PgV-only lysate) and PgVV (yellow). We considered the latent period finished when the number of free virions reached 1.3 than t0. n = 4. **b.** Virulence of PgV in mixed (pink) and solo (green) infections. Virulence was calculated as how many individual infections end in lysis. n = 1149 (PgV-only), n = 1824 (PgV/PgVV mix). **c.** Burst size of PgV in a mixed (pink) and solo (green) infections. n = 5. **d.** Cell survival (measured by chlorophyll A autofluorescence) of *P. globosa* in control and infected cultures. n = 5. **e.** Intracellular DNA copies (absolute copy numbers) of *P. globosa*, PgV and PgVV during infection. Three representative replicates are shown. A single outlier is marked with a circle in III. **f.** RNA copies (normalized to RNA concentration) of MCP and DNA polymerase of PgV and PgVV during infection. Three representative biological replicates are shown (same as in panel e.). Results for all six replicates can be found in Supplementary File 2.

The same experimental layout was followed to evaluate the effect of PgVV on the course of PgV’s infection. The latent period for both viruses was 8-9 hrs, regardless of PgVV presence (Fig. 2a, Supplementary File 2), a little shorter than previously reported^30^. We considered the latent period finished when free viral particles increased by 30% compared to the first sampled time point. We compared the virulence of PgV with and without PgVV, in our setup only ∼10% of infections ended in lysis, regardless of PgVV (Fig. 2b, Supplementary File 2), similarly to reports for other giant viruses^31–33^. In mixed lysates, successful co-infections of PgV and PgVV are rare: less than 20% of successful PgV infections are co-infections with PgVV (Supplementary File 2). A number of reasons could lead to this result. First, *P. globosa* defense mechanisms against viral infection might affect both viruses, while PgV might have additional anti-virophage defense systems. Second, it is likely that in contrast to Sputnik which enters the host-cell entangled in its host virus surface fibers^34, 35^, PgVV has to recognize, attach and enter the PgV-infected host independently. PgV particles lack long fibers (Fig. 1c) and we found no homologs of the fiber-associated proteins of mimiviruses^34, 36^. However, phage fiber-like proteins were found encoded in the PgV and PgVV genomes (see below). The timing of entrance of the virophage might be critical for a successful infection, for it depends on PgV for transcription, as we see synchronous expression of MCPs and DNA polymerases of both viruses (Fig. 2f). This is further supported by the presence of the early promoter motif of Mimiviruses in both PgV and PgVV (Fig. 3c).

**Figure 3.**
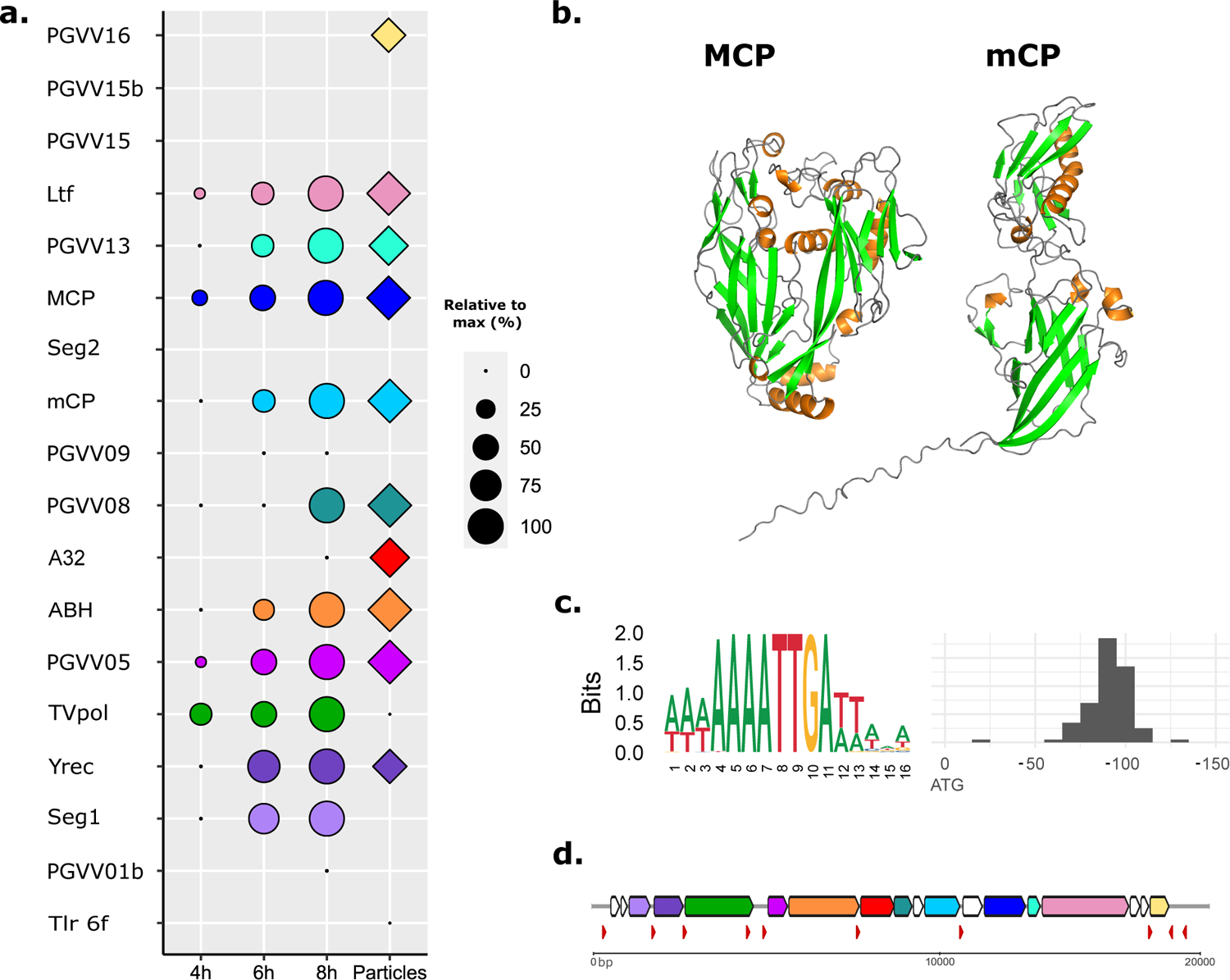
PgVV proteomic features. **a.** Proteins found by mass spectrometry in purified viral particles (rhomboids) and 4, 6 and 8 hours post-infection (circles). Relative quantification as described in the methods section. Proteins for whom peptides were found, yet below the significant threshold are marked with a point (0%). Tlr-6f, conserved uncharacterized protein; Seg1, GIY-YIG family nuclease; Yrec, OLV11-like tyrosine recombinase; TVpol, hybrid transposon-viral polymerase, superfamily 3 helicase; ABH, alpha-beta hydrolase (putative lipase); A32, ATPase; mCP, minor capsid protein; Seg2, homing endonuclease; MCP, major capsid protein; Ltf, L-shaped tail fiber-like protein; hypothetical proteins are denoted with their PgVV genomic serial number. **b.** Structural models of PgVV’s major (MCP) and penton (mCP) capsid proteins, featuring a double and a single jelly-roll respectively. Secondary structure elements are colored green for beta-strands, and orange for alpha-helixes. **c.** Early genes promoter motif of PgV-14T and motif location relative to the start codon of PgV genes (marked as 0). Numbering denotes bp. **d.** Schematic map of PgVV-14T genome, PgV early-genes promoter motif is marked with red arrows oriented according to their strand. ORFs color coding according to 3a, colored proteins were found in our proteomic data, white-colored ORFs were not significantly detected.

After inoculating the cultures with mixed and PgV-only lysates, we see immediate adsorption, and the number of free viruses remains low until 6-9 hrs post-infection (Supplementary File 2). *P. globosa*’s DNA was usually degraded during infection, probably due to early lysis of cells, however some replicates showed little host DNA degradation during the first hours (Fig. 2e, Supplementary File 2). PgV and PgVV genome copy numbers increased at 4 hours post-infection and kept increasing until burst (Fig. 2e). We selected two genes, representative of classical early (DNA polymerase) and late (MCP) genes, and measured their expression levels during the infection (Fig. 2f). PgV and PgVV DNApol transcripts can be traced already at 2 hours post-infection, while MCP transcripts appear after 4 hours. These results suggest that the virophage synchronizes its infection to that of its host virus, and supports our notion that the timing of entry is critical for PgVV’s success. Moreover, we see that the ratio between PgV and PgVV transcripts varies between experiments, even though the initial PgV/PgVV ratio was constant, possibly explaining part of the observed variability and the divergent PgVV burst sizes (Supplementary File 2).

The average burst size for PgV-only was 136 viruses cell^-1^ (minimum: 101; maximum: 181; s.d. = 28.29), lower than previously reported^30^. When co-infected with PgVV, we saw a small but significant decrease in the estimated average burst size of PgV: 136 (s.d. = 28.29) vs 108 (s.d. = 33.38) viruses cell^-1^ (Student’s T-test, p-value = 0.04) (Fig. 2c, Supplementary File 2). Thus, similarly to other virophages^22, 23, 25^, PgVV inhibits PgV reproduction. Since PgV and PgVV co-infections are rare, and given the high variability of this experimental setup, the actual burst size for PgV from a single co-infection is expected to be lower. Nevertheless, its effect might be negligible at the population level since lysates derived from a single co-infection showed roughly the same PgV progeny as lysates from a single infection by PgV alone (Supplementary File 2). We estimated the burst size for PgVV based on the rate of successful co-infections for each experiment and found very large variation, from 9 to 321 virophages cell^-1^ (Supplementary File 2). These values could partially explain the high variability we see in the *P. globosa*-PgV-PgVV system in our laboratory setting (Fig. 2e,f, Supplementary File 2). From *P. globosa*’s perspective, the cultures collapsed after 10 hours (for high PgV/cells ratio) or 20 hours (for lower ratio), regardless of PgVV presence, or the PgV/PgVV ratio (Fig. 2d, Supplementary File 2). Following previous findings on the *Cafeteria*-CroV-Mavirus system, where the virophage prevents CroV from lysing the entire host population (when CroV is added at low quantities)^37^, we reproduced similar conditions by infecting *P. globosa* cultures with a PgVV lysate where PgV was below the detection level of a standard PCR reaction. During the first four days there was no change in the viral numbers, or between infected and control cultures. At day five we saw a slight increase in both viruses’ DNA copy number; and two weeks after inoculation, PgV’s genome copy number quintuplicated, while PgVV’s triplicated, and the infected cultures were completely lysed as opposed to the stationary-phase controls (Supplementary File 2). Overall, we see that although PgVV replication is adverse to PgV at the single-cell level, the effect of PgVV is not significant at the population level in our laboratory setup. The variable dynamics regarding viral and virophage virulence might explain the coexistence of these players in the environment, as previously proposed for other host-virus-virophage systems^38^. Changes in the local ratios between the algal host, the giant virus and the virophage might result in a different short-term “winner”, maintaining the equilibrium between them on the global scale.

### Proteomics analyses

Proteomic analyses were performed on samples from 4, 6 and 8 hrs post-infection with a mixed lysate, uninfected *P. globosa* cultures and purified viral particles. Only 44 PgV proteins were detected in purified PgVV-14T particles, in contrast to 114 to 142 in the mixed samples, possibly as carry-over of the purification process, lending further support to the existence of independent PgVV virions (Supplementary File 4). Overall, the identification rate of viral proteins from both viruses improved with the course of the infection, with most viral proteins peaking at 8 hrs. Peptides for fifteen out of eighteen proteins predicted in PgVV were detected in our samples, six of them in all replicates of purified viral particles (Fig. 3a). Three proteins were detected in at least two samples, yet below the significant threshold (*pgvv01*, *pgvv01b, pgvv09*). Curiously, despite the lack of detectable signal in the proteomics data, transcripts from all of the remaining ORFs were amplified at 4 hrs post-infection (Extended Data Fig. 3). The presence of capsid proteins in the early stages of infection, despite MCP not being transcribed, can be explained by the high number of virophages adhering to or entering the cells. MCP, mCP, Ltf, the putative lipase and the proteins of unknown function PGVV05 and PGVV08 were consistently found in the viral particles, thus likely being components of the PgVV virion. The finding of Ltf in the particles is consistent with its predicted similarity to bacteriophage tail-like fiber proteins (Supplementary File 1, Extended Data Fig. 4), and further suggests that it may be involved in mediation of the recognition or attachment to the host cell, while the putative lipase (ABH) might mediate the virophage entry to the host cell, as shown for other small viruses^39^. The Tlr6f protein (widespread among PLVs, virophages, giant viruses and bacteriophages) was not detected in any of the infected samples despite it being expressed on the mRNA level, likely because of its low abundance; however a few peptides were detected in purified viral particles, below the significant threshold (Supplementary File 4). The packaging-ATPase and PGVV16 were detected only in some replicates of the viral particles, where identification of low abundance peptides is easier, yet they do not seem to be an integral part of the virions, similarly to PGVV13 and Yrec, since they are not consistently found in all viral particles replicates (Fig. 3a).

**Figure 4.**
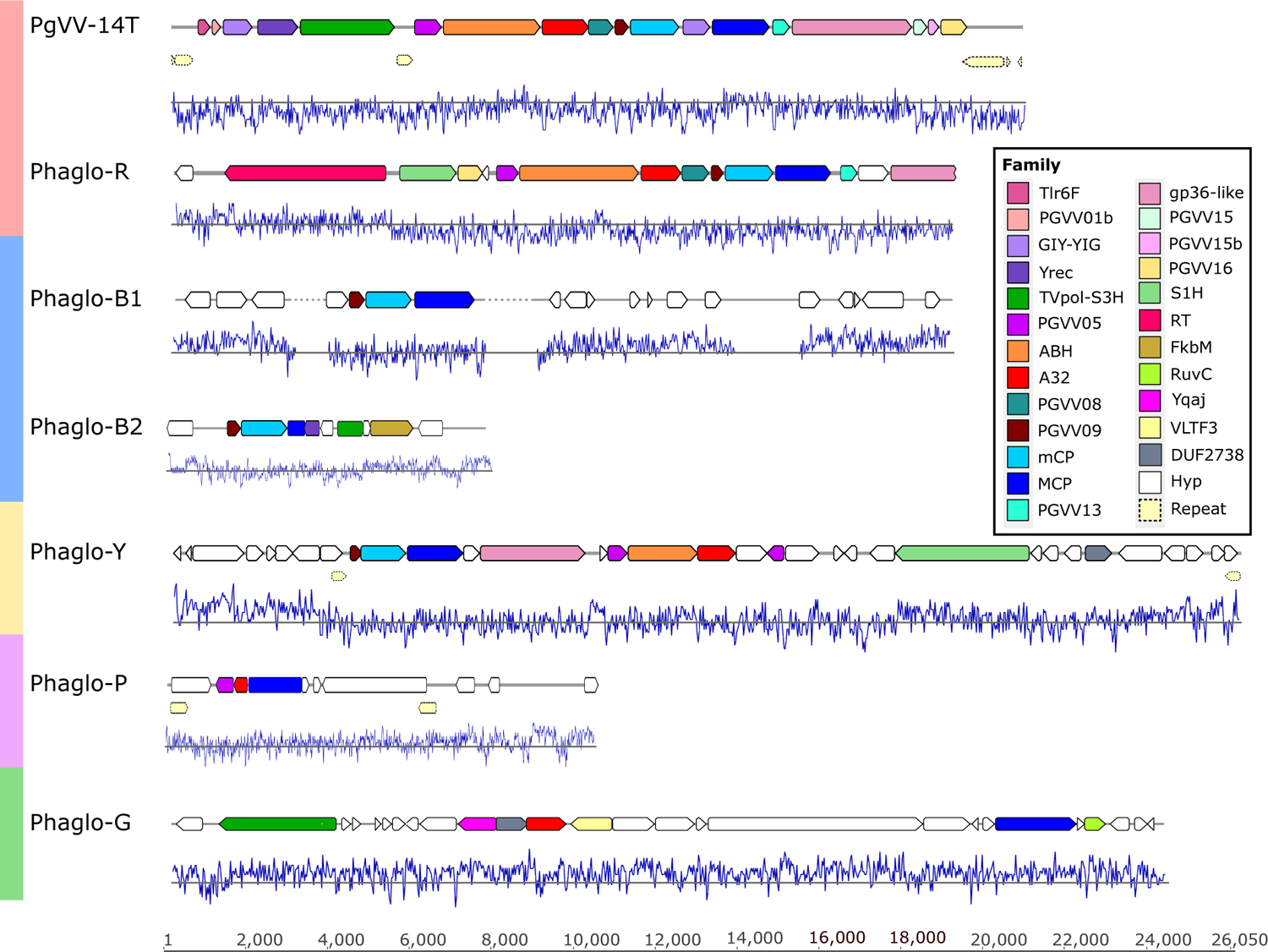
PLVs associated with *Phaeocystis globosa*. Schematic representation of the PgVV-14T virophage and *P. globosa* genomic contigs including viral-like elements. Gaps in the assemblies are marked with a dotted line. Homologous proteins are marked in color by their cluster family (Table S1), according to the legend. Repeats are marked in pale-yellow with dotted borders. Blue lines denote %GC content, the gray line marks 50% GC. Color-coding of the contigs names match the shading on Fig. 5a. The bottom bar designates bp. DNA sequences for the PLVs retrieved in this work can be found in Supplementary File 5. S1H; superfamily 1 Helicase; RT, reverse transcriptase; FkbM, methyltransferase; RuvC, RuvC nuclease; Yqaj, Yqaj-like recombinase; VLTF3, late transcription factor; DUF2738, unknown protein with DUF2738/5871 domain. The other designations are the same as in Fig. 3.

Regarding PgV, proteins containing extracellular domains PGV252, PGV055 and PGV041, are also present in the particles, probably related to attachment to the host (Supplementary File 4). A total of fifty-five proteins with no recognizable domains are found at least in 60% of the samples for viral particles, reminding us there is much more to these viruses than we currently know (Extended Data Fig. 5).

**Figure 5.**
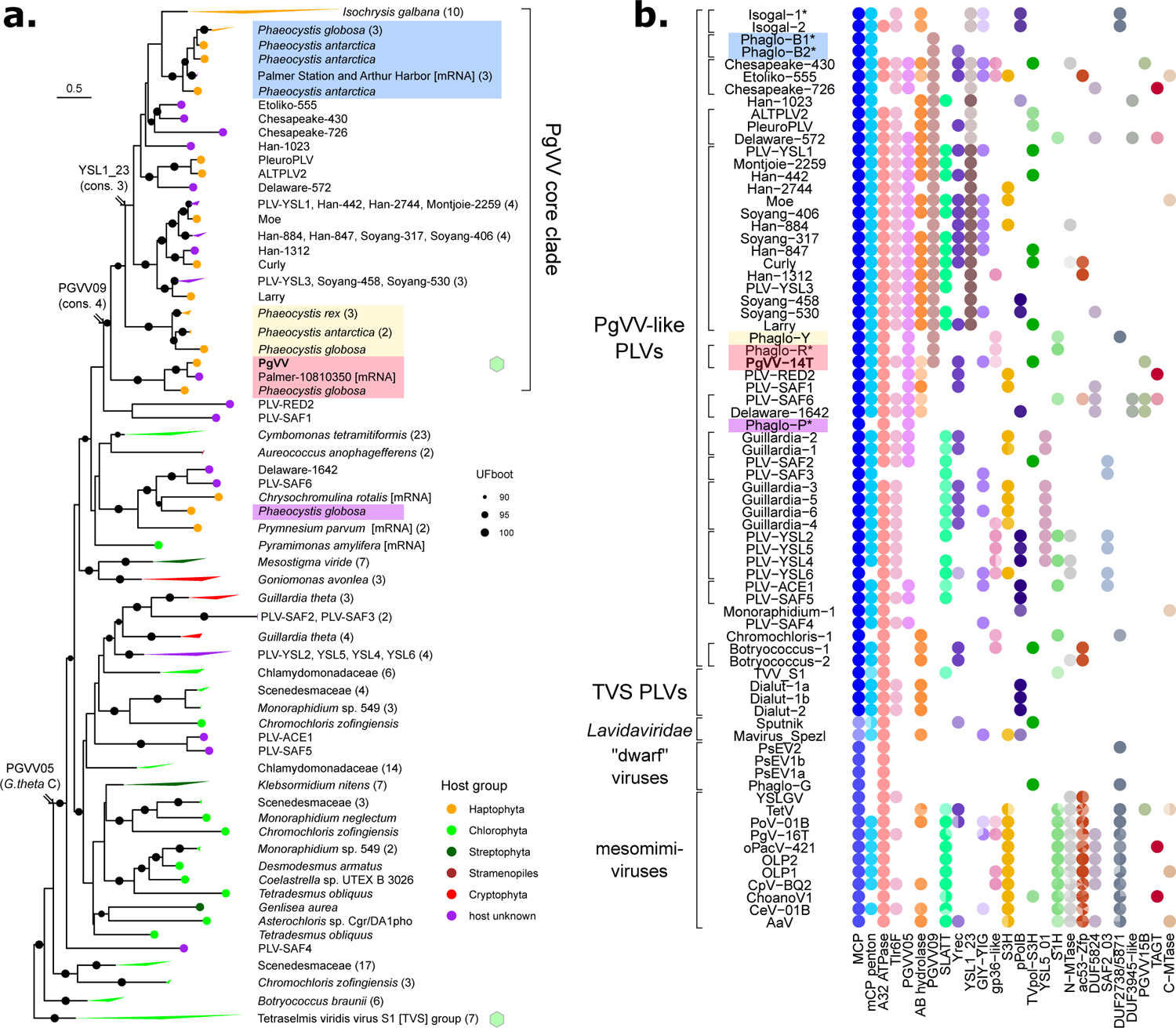
Phylogeny and gene content of PgVV-like PLVs. **a.** Phylogenetic analysis of MCPs from the PgVV-group PLVs. MCP sequences were extracted from previously published PgVV-group PLVs^8, 9, 19, 21^ and newly discovered viral elements found in eukaryotic genome assemblies. The tree is rooted by MCP sequences from the TVS group. Associated host groups are indicated when known. Four clades of PLVs discovered in *Phaeocystis* genome assemblies are highlighted with color (see Fig. 4). Parsimonious predictions of three gene acquisition events characteristic to the PgVV core clade are indicated. Cultured viruses, PgVV and TsV-N1, are indicated with a green hexagon. Numbers of sequences for collapsed clades are shown in parentheses. A complementary phylogenetic tree for the TVS group can be found elsewhere^46^. **b.** Core genes of PgVV-group PLVs. Protein sequences were clustered based on profile-profile matches and the clusters were further grouped into protein families based on shared Pfam matches. Each color hue represents one cluster, such that genomes may contain multiple clusters from the same family. Descriptions of the gene families are provided in Table S1. Only genes appearing in at least three subgroups of PgVV-type PLVs are shown, and are arranged by the number of genomes they are observed in. The genomes of the TvV-S1, virophages Sputnik and Mavirus, NDDVs and mesomimiviruses are provided for reference. Asterisks mark partial sequences. AaV – Aureococcus anophagefferens virus; CeV-01B – Chrysochromulina ericina virus CeV-01B; CpV-BQ2 – Chrysochromulina parva virus BQ2; PgV-16T - Phaeocystis globosa virus 16T; PoV-01B – Pyramimonas orientalis virus 01B; TetV - Tetraselmis virus 1; PsEV1a, PsEV1b and PsEV2 – *Pleurochrysis* sp. endemic viruses 1a, 1b and 2; TvV-S1 – Tetraselmis viridis virus S1. Species prefixes for integrated PLVs are as follows: Botryococcus – *Botryococcus braunii*; Chromochloris – *Chromochloris zofingiensis*; Dialut – *Diacronema lutheri*; Guillardia – *Guillardia theta*; Monoraphidium – *Monoraphidium neglectum*; Isogal – *Isochrysis galbana*; Phaant – *Phaeocystis antarctica*; Phaglo – *P. globosa*; Pharex – *P. rex*. Metagenome prefixes are as follows: ACE - Ace Lake; Chesapeake - Chesapeake Bay; Delaware - Delaware Bay, Etoliko - Etoliko Lagoon; Han - Han River, Montjoie – Lake Montjoie; RED – Red Sea; SAF – South Africa; Soyang – Lake Soyang; YSL – Yellowstone Lakes. For the complete list of viral genomes and all cluster and family assignments see Supplementary Files 5 and 6, respectively.

### PgVV depends on PgV for transcription

To get a better understanding of the procession of PgV infection, we divided its genes into early, middle and late, based on the proteomics results (Supplementary File 4). We assume that peptides detected in samples taken 4 hrs post-infection were transcribed as early genes, while peptides detected only at 8 hrs post-infection belong to late genes. Analysis of sequences upstream of the genes categorized as early yielded a conserved motif with the sequence AAAATTGA at its core (93 sites, E-value: 1.7e-61) (Fig. 3c). AAAATTGA was first reported as an early promoter motif in Mimivirus^40^ and related motifs appear to be common in other mesomimiviruses (Extended Data Fig. 6). Querying the PgVV genome with the early promoter motif brought 10 highly significant matches of which five fell upstream of start codons of *Yrec*, *TVpol*, *pgvv05*, *seg2* and *pgvv16* (Fig. 3d). The presence of the early promoter motif upstream of both the virophage and the virus DNA polymerase genes matches our observations on the co-expression of these genes (Fig. 2f). Interestingly, these results are in marked contrast to the Mimivirus-Sputnik and CroV-Mavirus systems where the virophage extensively uses the virus late promoter^16, 23, 41^. The presence of the early mimiviral promoter upstream of some of the ORFs and of AT-rich stretches in intergenic regions indicates that PgVV genes are expressed as monocistronic units, despite all of the ORFs having the same orientation in the genome (Fig. 4). Accordingly, we could not detect transcripts spanning adjacent gene pairs in our samples (Supplementary File 4), while individual genes could be amplified.

### *Phaeocystis* genomes contain PLVs related to PgVV

Given the close ties of PgVV to integrated algal PLVs^8^, we reasoned that closely related PLVs might be residing in the genome of *P. globosa*. To this end we created a partial assembly of the *P. globosa* genome^42^ and developed a bioinformatic pipeline based on co-occurrence of viral marker genes of PLV, virophage or NCLDV origin. Genomic fragments harboring at least two markers were further scrutinized for the presence of inverted repeats and drops in %GC content, as shown for integrated viruses in *Cafeteria sp.*^16^. Five scaffolds from *P. globosa* contain at least three PLV marker genes (Fig. 4). Based on phylogeny of the MCP we subdivided these fragments into four groups: Phaglo-R (shaded red) that clustered with PgVV, and the more distantly related Phaglo-Y (yellow), Phaglo-B1 and Phaglo-B2 (blue), and Phaglo-P (purple) (Fig. 5a). The blue and yellow groups are also present in the draft genome of *P. antarctica*, while the yellow group was additionally found in *P. rex*, suggestive of their distinct host range or time passed since their integration. Those contigs contain viral fragments representing PLVs of various degrees of completeness, with short flanking regions seemingly from the host. Similarly to the integrated Mavirus in *C. bukhardae*, *P. globosa*’s integrated PLVs have a lower %GC content than the rest of the host genome and the flanking regions on the scaffolds (37.5%-54.5% vs. 64.5%) (Fig. 4). An additional viral-like scaffold, Phaglo-G, has a high GC content (61.1%) and is flanked by short tandem repeats. MCP phylogeny and gene composition of Phaglo-G, found also in *P. rex*, revealed it to be related to *Pleurochrysis* sp. “endemic viruses”, and their origin is unclear (Extended Data Fig. 7). Given their size and affinity to NCLDVs we provisionally refer to these viruses as “NCLDV-like dwarf viruses” (NDDVs).

Neither transcripts or proteins encoded by *P. globosa* PLVs could be detected in our samples (Supplementary Files 4,5). It is possible that these PLVs respond to infection by prymnesioviruses other than PgV-14T. Indeed, we found transcripts for MCP similar to PgVV-like PLVs (Phaglo-R and Phaglo-B) in environmental metatranscriptomes from marine and polar environments (Fig. 5a, Supplementary File 5).

### PgVV-like PLVs belong to a distinct clade

Phylogenetic analysis of PLVs-MCP proteins from algal and protist genome assemblies revealed that most of the haptophyte-associated PLVs form a single well-supported clade, together with MCPs from freshwater and marine metagenomes, the “PgVV core clade” (Fig. 5a,b). A minority of haptophyte-associated MCPs, including Phaglo-P, form a second separate clade. Interestingly, MCP proteins from green algae are paraphyletic with respect to MCPs associated with haptophytes, cryptophytes and a stramenopile which indicates that the PgVV-like PLVs originated as green algal viruses (Fig. 5a). Integrated PgVV-like PLVs could be also found in *Isochrysis galbana,* while the genome of *Chrysochromulina parva*, the host of the PLVs Curly, Larry and Moe, did not possess any integrated viruses (Extended Data Fig. 8).

### PgVV-type PLVs Core Genes

To create a framework to uniformly classify virophages and PLVs, we performed a search for clusters of orthologous genes across PgVV-like PLVs. Some orthologous clusters could be further connected with each other based on the presence of a set of commonly conserved protein domains. Five PgVV proteins comprise the set of core genes present in nearly all PgVV-like PLVs, integrated and free alike: MCP, mCP, A32, PGVV05 and Tlr6F (Fig. 5b). The first three are also part of the core gene repertoire in TVS-like PLVs and the *Lavidaviridae* family of virophages (Fig. 5, Extended Data Fig. 9)^8,43^. The PGVV05 cluster includes the *G. theta* protein C^8^ and appears to be specific to the lineage of PgVV-like PLVs. The function of PGVV05 is unknown, but we hypothesize that they constitute a component of the virion (Fig. 3). The Tlr6F-like proteins are widespread beyond the clade of PgVV-like PLVs^8^, and in particular are present in PgV and other mesomimiviruses. Two further genes of unknown function appear to be restricted to the PgVV core clade: PGVV09 found in all members of the PgVV-core clade, and YSL1_23 that is absent from PgVV itself.

Two other widespread interesting gene families are ABH and Ltf. Sequence diversity and high variability of ABH active-site positions (ranging from GHSQGG in PgVV to SYSDGG in Phaglo-R) in these proteins is indicative of multiple independent acquisition events. Functions of alpha/beta-hydrolases are difficult to predict^44^, although based on the entry mechanism of other small viruses into their host cell^39^, and the ABH being consistently found in PgVV-14T capsids (Fig. 3), it is plausible that these proteins are lipases. *Ltf* is the longest ORF in PgVVs genomes and encodes a protein containing repeats with homology to gp36 of the T4-bacteriophage, a component of the long tail fibers (for more details, see Supplementary File 1). This protein was also consistently found in PgVV-14T viral particles, suggesting that it might be involved in mediating attachment to the host cell.

Tyrosine recombinase (Yrec) genes represent the most widespread family involved in genetic information processing among PgVV-like PLVs. Yrec is a feature shared between PLVs, Polintons and virophages and likely mediates their integration in host genomes^8^. GIY-YIG superfamily endonucleases have a more sporadic distribution. Surprisingly, PgVV-14T encodes two such endonucleases: Seg1 and Seg2, with Seg2 unusually located between the major and minor capsid proteins (Extended Data Fig. 10). Seg2 is likely an intronless, site-specific homing endonuclease, a selfish genetic element capable of integrating itself next to the recognition site (for more details see Supplementary File 1). This gene could serve as a defense system of PgVV against other similar virophages^45^, or be a SGE hitchhiking on PgVV.

### PgVV classification

Since the defining features of a *bona-fide* virus are the possession of a coat-protein encoding gene and the ability to form virions^47, 48^, we can now affirm that the Polinton-like PgVV-14T is a virus leading a virophage life-style. PgVV fulfills the basic requisites to be classified within the *Lavidaviridae* family of virophages: it depends on PgV for replication, and it encodes for (distant) homologs of the morphogenetic genes conserved in virophages^43^. However, based on phylogeny of core genes, PgVV is not directly related to the previously isolated virophages^8, 43^, thus we propose to classify PgVV separately. The core PgVV clade is well-supported in the MCP phylogeny, and also by association with haptophytes and the presence of the PGVV9-cluster genes. Based on those features, we propose to create a viral genus that would include PgVV-14T, PgVV-16T, PgVV-like PLVs integrated in haptophyte genomes, the putative *Chrysochromulina* virophages Larry, Curly and Moe^21^ and related PLV currently known only from environmental sequences. The NDDVs, including Phaglo-G and *Pleurochrisys* endemic viruses should be classified separately. Within this genus different species should be created to accommodate viruses infecting different hosts. Classification of metagenomically retrieved PLVs remains a challenge since very little is known regarding the possible life style and host range of these viruses. We suggest classifying the PgVV-like PLVs from *Phaeocystis* genomes together with PgVV-14T, until viral particles are isolated and characterized.

## Conclusions

Virophages, dsDNA viruses that parasitize an active virocell created by a NCLDV, have been known for more than ten years, yet only a few were successfully cultured. How these viral parasites shape and affect their host’s evolution remains to be studied for each particular system. Yet there are general features that seem to characterize the virophage lifestyle, like the ability to integrate to the cellular host genome, and the fitness cost to the viral host. In this work we isolated and characterized the Polinton-like virus PgVV-14T, a virophage infecting *Phaeocystis globosa* Pg-G(A). We show that PgVV-14T is a *bona-fide* virophage of linear dsDNA coated by a proteinaceous shell composed by major and minor capsid proteins, a putative lipase and proteins of unknown function. Based on the various related PLVs found in *P. globosa* it might also be capable of transiently or permanently integrating in its algal host genome. The fact that PgVV is an independent virophage confirms that integrated and free-standing PLVs carrying homologous capsid proteins could also form virions. Whether those PLVs are independent viruses like TsV-N1, have a virophage life-style, or replicate solely by a lysogenic/transposable life cycle remains to be studied. The existence of PgVV-14T viral particles sheds light on a poorly understood phase of viral evolution, for PgVV and its related PLVs resemble ancestors to dsDNA eukaryotic viruses. As virophages continue to be isolated and characterized, we get to glimpse into the work of the parasitism master, modulating the populations and evolution of viruses and cellular organisms, from the outside and within.

## Methods

### Cultures of *Phaeocystis globosa* and viruses

Non-axenic *Phaeocystis globosa* strain Pg-G (A), and the PgV-14T lysate from the NIOZ Culture Collection were used for this study. *P. globosa* was grown in Mix-TX medium (1:1 mix of f/2^49^ and ESAW^50^, enriched with Tris-HCl and Na_2_SeO ^50^), at 15°C and 90 µmol photon m^-2^ s^-1^ in a light/dark cycle of 16:8 hours. Experiments were conducted in exponentially growing cells. Large culture volumes (> 5L) were grown with gentle stirring on a magnetic stirrer. PgV and mixed PgV/PgVV lysates were obtained by inoculation of *P. globosa* cultures in late-exponential phase. After full lysis the lysates were gently filtered through a 0.45 µm filter (either 33 mm Millex SLHV033RS Millipore, or 75 mm Nalgene rapid flow filters – Thermo Fisher Scientific, depending on the lysate volume).

### Sequencing and assembly of PgV and PgVV

One ml of lysed *P. globosa* culture was filtered through a 0.45 µm filter and used to extract DNA using the Promega Wizard columns protocol^51^. Nextera libraries were sequenced using an Illumina MiSeq sequencer at the Technion Genome Center, Israel. The raw data was de-replicated with ParDRe v. 2.1.5^52^ and trimmed with trim_galore v. 0.6.6^53^. The genome assembly was performed with spades v. 3.14.1^54^. Additional Sanger reads were generated to close assembly gaps in the PgV-14T genome (see Supplementary File 2 for primers list). PCR was performed with Ex-Taq enzyme (TaKaRa) in a total volume of 30 µl containing 1 µl viral DNA, Ex-Taq buffer (×1), 0.8 mM primers, 0.8 mM dNTPs and 0.75 U polymerase. PCR conditions were as follows: 95°C – 5 min, 30 cycles of 95°C – 30 sec, 60°C – 30 sec, 72°C – 5 min, and a final elongation of 72°C – 5 min. PCR products were cleaned from gel using NucleoSpin Gel and PCR cleanup (MN) and cloned in TOPO-TA plasmids (Invitrogen) according to manufacturer’s specifications. Sanger sequencing was performed by Macrogen Europe.

The terminal inverted repeats of the PgVV-14T were represented as separate fragments in the spades assembly and thus the following strategy was utilized. The PgVV scaffold was trimmed to include only the non-repeated region and extended with ContigExtender using the raw data^55^. The scaffold was trimmed to include a minimal region that would contain the fragments. Given the high sequence similarity between PgV and PgVV isolates, ORFs could be directly transferred from PgV-16T and PgVV-16T to PgV-14T and PgVV-14T, respectively.

### Electron Microscopy

For transmission electron microscopy (TEM) 20 L of exponentially growing (late-stage) *P. globosa* were infected with a mix of PgV-14T and PgVV-14T viruses. Upon full lysis the lysate was filtered through 0.45 µm filters (Nalgene rapid flow filters – Thermo Fisher Scientific) to remove cell debris. The filtrate was concentrated using a 100 kDa TFF column (Repligen N06-E100-05-N) and viruses were pelleted by ultracentrifugation (141,000 × g, 2 hrs, 4°C). The viral pellet was resuspended in Mix-TX medium, loaded into an Optiprep (Sigma) 25-40% stepped gradient, and centrifuged at 160,000 × g (SW 41-Ti rotor), for 15 hrs, 4°C. Bands were pulled using a syringe to a Millipore Amicon ultra 100,000 K (Mercury), and centrifuged several times at 5,000 × g to change the medium back to Mix-TX. Ten µl samples were loaded into grids and stained with 10 µl 1% uranyl acetate for 1 min, followed by air-dry desiccation (3 hrs). Transmission electron microscopy was performed in a Talos L120C transmission electron microscope at an accelerating voltage of 120 kVe at Rappaport Faculty of Medicine, Technion. Particle sizes were calculated for both viruses using ImageJ v.1.53q^56^.

### Identification of PgVV particles by PCR and SYBR staining

To verify the existence of PgVV virions, 0.45 µm filtered viral lysates (6 ml) were separated into three fractions: One fraction remained untouched (L – Lysate), the two other fractions were filtered twice through 0.2 µm filters (Millex Syringe-driven filter units, SLGV033RS, Millipore) (F – filtered), one of them was then boiled for 10 minutes (B – Boiled). DNAse (Ambion Turbo DNAse cat. AM1907) was added to all three fractions in a 50 µl reaction, as follows: buffer (x1), 2U DNAse, 44 ul sample, and incubated 30 min at 37°C. After 30 min, additional 1U of DNAse was added to each sample and further incubated for 30 min at 37°C. DNAse inactivation was performed according to manufacturer instructions. PCR was then performed on all three fractions for PgV and PgVV marker genes with primers 7,8 (MCP) and 9,10 (TVpol) respectively (Supplementary File 2). PCR was performed using the Bio-Ready Mix 2X colored (Bio-Lab) with the following parameters: 95°C – 5 min, 30 cycles of 95°C – 30 sec, 60°C – 30 sec, 72°C – 30 sec, and a final elongation of 72°C – 5 min.

Since PgVV genome was found to be more abundant than PgV’s (also reported in^27^), SYBR-stained lysates were prepared for visualization. Viral lysates (PgV-only, PgVV-only and Mix) were filtered through 0.45 µm filters, and stained with SYBR Green-I as described elsewhere^57^, and manually counted in an Elyra 7 eLS microscope at the Technion LS&E with the Plan-Apochromat 63x/1.4 Oil DIC M27 objective, a Scientific Scmos camera and the 1.4-420782 lens. Images were taken with a 488 nm excitation wavelength and 515 nm emission wavelength for 100 ms, using a FITC 525/50 filter and rendered using the SIM^2^ algorithm. Particle analysis for three to four field views was performed with 3D Objects Counter v.2.0.1 for ImageJ v.1.53q^58^ with default settings. To make the analysis as unified as possible, a single value for thresholding was used for most of the PgV and Mix fields. Due to the presence of a significantly brighter background and of spots that appeared to be damaged viral particles in the PgVV-only images, for PgVV analysis a separate higher threshold was chosen.

### *P. globosa* and PgV counts

*P. globosa* cells were counted using flow cytometry on the basis of their scattering (SSC) and autofluorescence using a 488 nm air-cooled blue laser (530/30 BP filter and 505 LP filter), with a BD-LSRII flow cytometer. PgV particles were counted in a BD-LSRII flow cytometer and in the Cytek Aurora Flow Cytometer after fixation and staining by SYBR Green-I, based on FSC and the 530/30 BP, 505 LP laser, as described elsewhere^59^. Flow cytometry was used to count cell and PgV abundance when necessary, however, PgVV could not be detected using this approach. Therefore, in experiments where we compare PgVV to *P. globosa* and/or PgV abundances, we quantified their DNA copies using qPCR (as described below). This enabled us to have a uniform (yet inflated) estimation for each entity that allows numerical comparisons. In all cases where qPCR was used to estimate PgV DNA copy numbers, the samples were also counted by flow cytometry to confirm that we observe the same pattern in the experiment using both approaches. Additionally, *P. globosa* cultures were monitored using chlorophyll A auto-fluorescence as a proxy for bulk biomass and livelihood of the cells (excitation/emission: 440/680 nm) in a Synergy 2 microplate reader (Bio Tek) in experiments where exact cell number was not required, as the flow cytometers were not always available for use.

### Isolation of a pure PgV lysate

A mixed lysate containing PgV and PgVV was diluted by 5×10^-5^ to ensure one viral particle per 10 µl and used to infect 380 aliquots of *P. globosa* cultures in 96-well plates (200 µl) and incubated for 10 days. Lysed cultures were checked for PgVV presence by PCR.

### PgV and PgVV growth curve experiments

To measure the latent period of PgV and PgVV we used a PgV/*P. globosa* ratio of 0.1 (counted by FACS), and a PgV/PgVV ratio of 1 (calculated by qPCR). Experiments were performed in 25 ml of *P. globosa* cultures. At every sampling point (0, 2, 4, 6, 8, 9 and 10 hrs post infection) 2 ml culture was filtered through a 0.45 µm filter (Millex, SLHV033RS, Millipore) and the filtrate was kept at 4°C until analysis (for a maximum of 2 weeks). Samples were treated with DNAse (as described above), DNA was extracted using the Promega Wizard columns as described elsewhere^51^, and quantified by qPCR (as described below). PgV fixed particles were also counted by Cytek Aurora Flow Cytometer as described elsewhere^59^. We considered the latent phase finished when the free viruses reached 1.3 times the free viruses at 0 hrs. n = 5. Experimental results for this and all the experiments described below can be found in Supplementary File 2..

### PgV and PgVV viral progeny calculation

To assess population-level fitness costs of PgVV infection on PgV, we used an estimate of PgV progeny obtained as the yield of viral particles from a single infection of a *P. globosa* cell in a 200 µl culture volume. 96-well plates with 200 µl of exponentially growing *P. globosa* cultures were infected with a diluted PgV-only or PgV/ PgVV lysate, such that every well will be inoculated with a maximum of one PgV particle. After lysis PgV particles were counted by flow cytometry while PgVV presence was confirmed by PCR. n = 3, a total of 14 lysed wells were analyzed.

### PgV and PgVV burst size calculation

Burst size calculations were performed in exponentially growing *P. globosa* cultures (5 ml) infected with a virus/cell ratio of 10-50. Lysates and cultures were counted before the experiment and after full lysis, *P. globosa* and PgV were counted by flow cytometry as described above, while PgV and PgVV DNA copy numbers were counted by qPCR as described below. Burst size of PgV was calculated by subtracting the original number of viruses from the final count (as counted by FACS) and dividing the difference by the number of cells in the original culture. A proxy for PgVV burst size was calculated by subtracting the original PgVV number from the final PgVV number (calculated by qPCR), and then dividing the difference by the calculated number of successful PgV infections from the same experiments (as calculated in Supplementary File 2). n = 5.

### Virulence of PgV

To calculate virulence (proportion of infections ending in lysis), exponentially growing *P. globosa* cultures were infected with PgV only and a mix of PgV and PgVV at a virus/cell ratio of 30-50. Since adsorption of viruses was very fast (Supplementary File 2), and at 6 hrs we already see lysis of the cells, we chose 3 hrs post-infection as the time point for analysis as it gives enough time for viral adsorption, yet short enough to have intact cells by the end of the sorting. Three hours after infection the cells were pelleted at 5,000 × g for 3 minutes and washed with fresh media three times to remove all free viral particles. Infected *P. globosa* cells were sorted into 96-well plates of fresh exponentially growing *P. globosa* cells using the FACSAria III sorter as described elsewhere^60^ and incubated until full lysis. PgVV presence in lysates was confirmed by PCR. 1149 wells were surveyed for PgVV-only, 1824 for PgV+PgVV. An uninfected culture of *P. globosa* was subjected to the same treatment as a control for the livelihood of the cells.

### *P. globosa* cell survival

Cell survival experiments were conducted on exponentially growing *P. globosa* cultures infected with either PgV only or a mix of PgV and PgVV at a virus/host ratio of 3-10 and incubated for two days. Cell survival was measured by chlorophyll A auto-fluorescence as a proxy for bulk biomass and livelihood of the cells (excitation/emission: 440/680 nm) in a Synergy 2 microplate reader (Bio Tek). n = 5

### PgVV-only infection of *P. globosa*

Exponentially growing *P. globosa* cultures were split each into four 200 µl cultures: Uninfected control, infected with PgV only, infected with PgVV only and infected with a mix of both. A PgVV-only lysate was obtained by 0.2 µm filtering of a mixed lysate (Millex, SLGV033RS, Merck Millipore), TFF concentration (Repligen N06-E100-05-N, 100 kDa) optiprep (Sigma) gradient separation (as described above) and filtration through a 0.1 µm filter (Millex, SLVV033RS, Merck Millipore). PCR of the resulting lysate showed PgV below the detection level for 30 cycles. The mixed lysate was obtained by combining the PgV-only and PgVV-only lysates. Cultures were infected with lysates in a 20% v/v ratio (ensuring at least 3 viruses per cell) and incubated until the control culture declined (4 days after infection). Growth of *P. globosa* was monitored by measuring chlorophyll A OD (excitation/emission: 440/680 nm) in a Synergy 2 microplate reader (Bio Tek). Samples from the infected cultures were diluted in TE and kept at −20°C until analysis (maximum of 2 weeks). Samples were further diluted in DDW (final dilution 1:100) and analyzed by real-time qPCR as described below. The same setup was used for a PgVV/PgV co-infection at high virophage/virus ratio (proxy of 20 virophages per giant virus, calculated using qPCR copies, yet PgV was below detection for a standard PCR reaction of 30 cycles). Cultures were incubated for two weeks. n = 3 for each experiment.

### Course of PgV and PgVV infection

Infection experiments were performed to obtain intracellular DNA, RNA and proteins during the course of an infection cycle. Exponentially growing *P. globosa* cells were infected with a PgV/PgVV mixed lysate, at virus/host ratio of 3-10. For intracellular DNA 1.5 ml culture was pelleted by centrifugation at 10,000 × g, 10 °C for 7 min, and resuspended in 2 ml media, three times. Washed pellets were flash-frozen and kept at −80°C until DNA extraction (maximum of 2 months). DNA was extracted using the GenElute – Plant Genomic DNA Miniprep Kit (Sigma), samples were cleaned by DNAse (as described above), and analyzed by qPCR as described below. For RNA extraction 1.5 ml culture was pelleted by centrifugation (7 min, 10,000 × g, 10°C), flash-frozen and kept at −80°C (maximum of 2 months). RNA extraction was performed with the Monarch Total RNA Miniprep Kit (NEB). DNAse treatment was performed as described above and cDNA was synthesized using LunaScript RT Supermix Kit (NEB). RT and non-RT vials were checked by PCR with primers 7,8 for PgV (Supplementary File 2). cDNA was diluted by ×100 and used as template for qPCR. 50 ml of culture was pelleted for proteomic analyses by 20 min centrifugation at 5,000 × g, followed by another 5 min at 10,000 × g. Samples were flash-frozen and kept at −80°C (for up to 12 months). The *P. globosa*-PgV-PgVV system showed high variability between biological replicates, especially regarding relative RNA amounts. In experiments where averaging replicates resulted in a graph that does not represent the diverging trends of the experiments, we present one experiment per trend separately (Fig. 2). n = 5

### Real-Time qPCR

qPCR reactions were performed on extracted DNA & RNA, and on diluted lysates according to the description for each experiment (see above). The PerfeCTa SYBR Green Fast Mix (QuantaBio) was used in a volume of 20 µl: 5 µl template, MasterMix ×1 and 0.25 mM primers. Reactions were carried out on a LightCycler 480 Real-Time PCR system (Roche) as follows: 95°C – 10 min, 45 cycles of 95°C – 10 sec, 60°C – 30 sec (annealing and elongation). At the end of each cycle the plate fluorescence was read and the point at which the fluorescence of a well raised above the background was calculated using the LightCycler 480 software (release 1.5.0) using the absolute quantification/second-derivative maximum analysis package. Specificity of each amplified qPCR product was verified by melting curve analysis on the LightCycler 480 instrument. DNA and RNA concentration of each sample for every single experiment were measured using Qubit (dsDNA and RNA high sensitivity kits - Thermo Fisher Scientific) and equalized to enable reliable and uniform quantification. In addition, standard curves were generated for each qPCR run from known copy numbers of a linearized pGEM-T Easy plasmid (Promega) containing the amplified PCR product to calculate the number of absolute copies in the samples.

### Proteomics sample preparation

Proteome analysis was performed to free purified viral particles, infected and control cells. For viral particles three independent *P. globosa* cultures (20L, 20L and 4L) were grown until late exponential phase and infected with three independent PgV and PgVV mixed lysates. An extra lysate of 10L was filtered through 0.45 µm and 0.2 µm (x2) to obtain a PgVV pure sample. Samples were prepared following the protocol described above for electron microscopy lysates preparation. For infection experiments, three time points were used for proteomic analysis, representing the early (4 hrs), middle (6 hrs) and late (8 hrs) stages of infection. For a preliminary experiment one replicate at 6 hrs post-infection was fractionated in a gel and seven fractions were run and analyzed separately, while a viral lysate (4L) was cut into three fractions from the gel. All other samples were analyzed whole, since the fractionation led to little improvement in resolution. Overall, three purified mixed viral lysates, one PgVV only lysate, three replicates for infected cells at each time point and one replicate for each time point of control cells were run in the mass spectrometer.

Frozen cells were resuspended with Tris-HCL pH 7.4 with a final concentration of 50 mM and 5% SDS and incubated at RT for 30-60 minutes. Cells were disrupted by beat beating for 2 minutes using 0.4-0.6 mm glass beads (Sartorius). Beads were pelleted by centrifugation (21,000 × g, 5 min, RT). Sonication of the samples was performed with a UP200St ultrasonic processor connected to vialtweeter (Hielscher, Germany) using a cycle of 80% with amplitude of 100% for 10 min. The samples were centrifuged (21,000 × g, 10 min, RT) and 90% of the sample volume was recovered. For viral particles, 1 ml of purified viral particles were mixed with Tris-HCl pH 7.4 to a final concentration of 50 mM and 5% SDS, and incubated 60 min at RT. Protein SDS PAGE-sample buffer with 10 mM β-mercaptoethanol was added to 10 ug protein (measured by Bradford Ultra) of viral lysates, control and infection samples, and incubated overnight at RT. Samples were run in a GeneScript gel (4-20%). Purified viral lysates were prepared and analyzed following in-gel digestion as described before^61^. Infected and control samples were digested in-solution using S-TRAP™ (Protifi, USA)^62^. Seventy five ug of each lysate were digested with sequencing-grade trypsin (Promega) 1:100 ratio for 12 hours at 37°C. Resulting peptides were desalted using C18 StageTips^63^ or TopTip™ (PolyLC, USA) and eluted with 50 μL of 50% acetonitrile, 0.1% FA, dried to completeness and resuspended in 2% acetonitrile, 0.1% FA.

### LC-MS

Desalted peptides of the different samples were subjected to LC-MS/MS analysis using Q-Exacitive-Plus or Q-Exactive HF mass spectrometer (Thermo Fisher Scientific) coupled to nano HPLC. The peptides were resolved by reverse-phase chromatography on 0.075 × 180 mm fused silica capillaries (J&W) packed with Reprosil reversed-phase material (*Dr. Maisch; GmbH, Germany*). The peptides of in-gel digested samples (purified viruses) were eluted with a linear 60 min gradient of 5–28%, followed by a 15 min gradient of 28–95%, and a 10 min wash at 95% acetonitrile with 0.1% formic acid in water (at flow rates of 0.15 μl/min). The peptides of in-solution digestion (infected and control samples) were eluted with a linear 120 min gradient of 6–30%, followed by a 15 min gradient of 30–95%, and a 15 min wash at 95% acetonitrile with 0.1% formic acid in water (at flow rates of 0.15 μl/min). Mass spectrometry analysis by Q Exactive Plus mass spectrometer (Thermo Fisher Scientific) was in positive mode using a range of m/z 300–1800, MS1 resolution 70,000 with AGC target: 3E6; maximum IT: 20 ms. These were followed by high energy collisional dissociation (HCD) of the 10 most dominant ions selected from the first MS scan. MS2 scans were done at 17,500 resolution, AGC target 1E5, maximum IT: 100msec, isolation window: 1.4 m/z; and HCD Collision Energy: 25%. Dynamic exclusion was set to 20 seconds and the “exclude isotopes” option was activated. Mass spectrometry analysis by Q Exactive HF mass spectrometer (Thermo Fisher Scientific) was in positive mode using a range of m/z 300–1800, MS1 resolution 120,000 with AGC target: 3E6; maximum IT: 20 ms. These were followed by high energy collisional dissociation (HCD) of the 20 most dominant ions selected from the first MS scan. MS2 scans were done at 15,000 resultion, AGC target 1E5, maximum IT: 60msec, isolation window: 1.3 m/z; and HCD Collision Energy: 27%. Dynamic exclusion was set to 20 seconds and the “exclude isotopes” option was activated.

### Proteomics Data analysis

Infected cultures and purified viral particles samples MS/MS spectra were analyzed with MSFragger version 3.5^64^, via FragPipe version 18.0 (https://fragpipe.nesvilab.org/) while using IonQuant (version 1.8) and Philosopher (version 4.3). The searches were conducted using Fragpipe LFQ-MBR configuration. Precursor mass tolerance was set to 20 ppm, fragment mass tolerance was set to 20 ppm, cleavage type set to “Enzymatic”, the enzyme was defined as strict trypsin and 2 missed cleavages were allowed. Cysteine carbamidomethylation was set as fixed modification and methionine oxidation and protein N-terminal acetylation were set as variable modifications. Peptide length was set to be between 7 to 50 amino acids and using default settings for label-free quantification. The searches were conducted against a database composed of viral proteins (PgVV, PgV), proteins encoded in the *P. globosa* integrated PLVs and *P. globosa* proteins. The set of proteins representing *P. globosa* was created by combining amino acid sequences encoded in the transcriptomes of three closely related strains: RCC851 (NCBI TSA HBRH00000000), RCC678 (HBRF00000000) and RCC739 (HBRB00000000). ORFs were predicted with TransDecoder v. 5.5.0 (https://github.com/TransDecoder) and the protein sequences were clustered at 100% identity level with cdhit v. 4.8.1^65^. Orthogroups were identified with ProteinOrtho v. 6.0.25^66^ using DIAMOND v.2.0.6.144^67^, identity threshold of 90% and coverage threshold of 25%. Representative proteins were selected from orthogroups appearing in at least two of the three strains.

For relative analysis of the infection course-proteomics we calculated and combined Max-LFQ intensity for all peptides found for a single protein at each time point. The time point with the highest score was arbitrarily set as 100% and the other two time points were normalized accordingly. For viral particles we used a yes/no approach, so that the 100% represents proteins found in the 5 samples run in the mass spectrometer, 80% for proteins found in 4 samples, etc. We analyzed only proteins with a Max-LFQ score, yet we also present results for proteins which peptides were detected below the significant threshold (Supplementary Figure 2).

### Modeling of PgVV capsid proteins

PgVV mCP (penton) and MCP proteins were folded with alphafold2 via ColabFold v.1.3.0.^68, 69^.

### Genome assembly of *Phaeocystis* species

In order to extract genes of viral origin and eventually complete viral segments from the genomes of *Phaeocystis* species we assembled the raw data available for genomes of *P. globosa* Pg-G (Bioproject PRJNA265550), *P. antarctica* CCMP1374 (PRJNA34537)^42^ and *P. rex* CCMP2000 (PRJNA534927), distributed publicly via NCBI SRA. Mate-pair libraries of *P. globosa* and *P. antarctica* were processed by detecting and replacing the junction linker with cutadapt v. 4.1 and custom scripts and categorizing reads with nxtrim v. 0.4.3^70^. All reads were trimmed trim_galore v. 0.6.7 (https://www.bioinformatics.babraham.ac.uk/projects/trim_galore/). The trimmed data were assembled with megahit v. 1.29^71^ using default settings. To extract complete viruses integrated in the *P. globosa* genome, the megahit assembly was scaffolded with SoapDenovo v. 2.40^72^ with K-mer size of 127 and remaining scaffold overaps were joined by a round of assembly with mira v. 5rc2 (clustering, accurate)^73^. Long viral segments of at least 6000 bp were extracted by searching for MCP genes with the HMM profile for PLVs and viropages^9^ and the NCLDV-specific profile VOG01840 from VOGDB (http://vogdb.org/). The extracted scaffolds were extended with Contig Extender v. 0.1^55^ and then polished and gap-filled with pilon v. 1.24^74^.

### Identification of MCP genes in assemblies

PgVV-type major capsid protein (MCP) genes were searched for in a collection of eukaryotic genomes and custom set of transcriptomes by extracting ORFs and searching using hmmsearch from HMMER v. 3.3.2^75^ with a HMM profile based on an alignment of MCPs from PgVV and *Phaeocystis* endognized PLVs with an E-value threshold of 1e-8 and sequences at least 200 amino acid residues were extracted for downstream analyses. The same analysis was implemented for NCLDV-type MCPs: initial search was performed with a HMM profile built from MCP sequences from mesomimiviruses and pre-extracted MCP sequences from endemic viral elements. The collection covered 186 species (see Supplementary File 5) with the four largest groups represented by green plants (96 species), Stramenopiles (46), Alveolata (11) and Haptophyta (10).

### Phylogenetic analysis

The collected MCP protein sequences were combined with MCPs from previously reported PgVV-group PLVs and TVS-group PLVs for rooting and aligned with hmmalign from HMMER using the virophage and PLV MCP profile from^9^. For the NCLDV MCPs the reference set included a representative set of MCP genes from *Mimiviridae* and *Phycodnaviridae* with *Iridoviridae*, *Ascoviridae*, *Marseilliviridae* and *Asfarviridae* and the alignment was performed with a HMM profile created from a mafft alignment of reference NCLDV MCPs. Alignments were trimmed to include aligned positions and resulting sequences longer than 300 residues after trimming were selected for phylogenetic analysis using iqtree v.2.1.2 with 1000 ultrafast bootstrap iterations^76, 77^. Shorter sequences were placed on the resulting trees by evaluating the trees with ng-raxml v.1.0.1^78^ and performing phylogenetic placement with epa-ng v.0.3.8.

### Identification of *Phaeocystis* PLVs in transcriptome data

PgVV-14T MCP protein sequence was used as query for BLAST against a collection of JGI freshwater and marine metatranscriptomes (updated by August 2021). Only hits to publicly available unrestricted databases were used.

### Gene homology and functional annotation

Homology between genes encoded by PgVV and related integrated and free PLVs was established by using profile-profile matches. All predicted protein sequences from PgVV-type PLVs, unrelated reference PLVs and virophages, integrated NCLDV-like dwarf viruses (NDDVs) viruses and mesomimiviruses were merged together and clustered with mmseqs v. 13.45111^79^ at a minimum of 30% identity and 80% coverage and the clustered sequences were aligned with result2msa. Each sequence was searched against the resulting database with hhblits from HH-utils v. 3.3.0^80^ (three iterations, E-value threshold of 1e-5) and secondary structure was predicted with addss.pl. The a3m database obtained this way was searched against itself with hhsearch from HH-utils. The results of the hhsearch matches were filtered to include hits with probabilities of at least 90 and a coverage of at least 60% of the query and the template and the resulting match pairs were clustered with MCL v. 14.137^81^. The same a3m database was used to query a profile database based on Pfam v. 34.0 available from the HH-suite webserver with hhsearch. Manually curated rules based on Pfam profile matches as well previously published annotations for individual genes were created to assign functions to the resulting clusters.

### Analysis of PGVV14 sequence

Protein sequence coded by *pgvv14* (*Ltf*) was analyzed by searching it against PDB_mmCIF70_21_Mar, Pfam-A_v35 and UniProt-SwissProt-viral70_3_Nov_2021 databases with hhsearch via the HHpred Server^82^ and by searching for repeats with RADAR via the EBI tools server^83^. PGCG_00042 and other PLV-encoded proteins matching the Pfam profile of T4 tail-fiber protein gp36 in local hhsearch (see above) with a probability as low as 80 were classified as proteins containing gp36-like domains.

### Promoter motifs in PgV

PgV genes were classified as early, middle or late genes based on their proteomic profile. Genes whose peptides were detected at 4 hours were considered “early”, while genes with detected peptides only in 8 hours post-infection samples were labeled “late”. The rest, genes whose detected peptide abundance did not change between the samples at 6 and 8 hours post-infection samples were considered “middle”. Many genes had no peptides detected and were classified as “none”. Although this division is not expected to mirror the exact RNA expression pattern of the genes (for example the MCP protein is detected at 4 hrs, yet it’s RNA is only starting to be transcribed), we expected the majority of genes classified as “early” to be in this category, and enable better resolution for the motifs analysis. 150 bp upstream from the starting codon of each gene were extracted and every category (early, middle and late) was analyzed separately using meme from the MEME suite v. 5.3.0 for a promoter motif.

Promoter motifs were searched across mesomimiviruses with meme from the MEME suite v. 5.3.0. Up to 10 promoter motifs with a length of 6-16 nt were searched for in the upstream 150 nt of each ORF. Sequences similar to the AAAATTGA-containing putative early promoter motif of PgV were searched in the whole genome of PgVV-14T using fimo from the MEME package using the meme output as query. The matches were filtered to include hits with q values < 0.05 and required the matched sequence to include the highly conserved TG nucleotide. For middle and late genes we could not detect a significant promoter motif.

## Acknowledgments

We thank Anna Noordeloos for providing advice on how to culture *P. globosa* and PgV, Lihi Shaulov for expert technical assistance with TEM sample preparations and imaging, Irena Pekarsky and Nitzan Dahan for their help with light microscopy, Ilana Navon and the Smoler Proteomics Center for their help with the mass spectrometry analyses, the ICTV Virophage study group for nomenclature discussions, and Shirley Larom for technical assistance. This work was funded by a European Commission ERC Advanced Grant 321647 (O.B.), Israel Science Foundation grants 143/18 (O.B.), 1623/17 and 2167/17 (T.L. and O.K.), and the Ariane de Rothschild Women Doctoral Program (S.R.). O.B. holds the Louis and Lyra Richmond Chair in Life Sciences.

## Author contributions

S.R. conceived the project and designed the experiments. S.R. performed all viral work. A.R. performed bioinformatic analyses. S.R., T.L. and O.K. performed proteomics. C.P.D.B supplied the algal and viral strains. O.B. supervised the project. S.R. wrote the paper, which was critically revised and approved by all authors.

## Data and code availability

The sequencing data are available from NCBI SRA SRR20333090 (Bioproject PRJNA835735). PgV-14T and PgVV-14T genome assemblies were deposited in NCBI Genbank under accession numbers OP080611 and OP080612. Annotated fragments of complete PgVV-group PLVs from *P. globosa*, *P. antarctica* and *Isochrysis galbana* genomes are provided as Supplementary File 7. Code used for bioinformatic analyses is available from https://github.com/BejaLab/PgVV. Source data are provided with this paper. Supplementary Files 2-7 will be provided upon request.

---

**Extended Data Figure 1.**
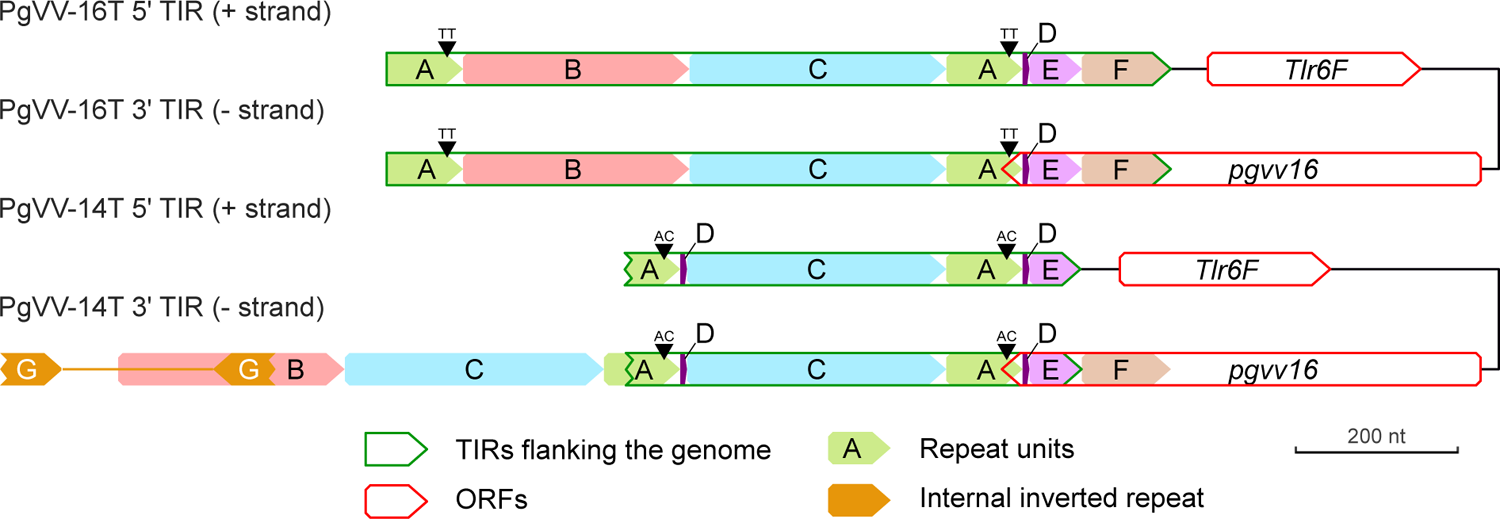
Comparison of the sequences of the terminal inverted repeats (TIRs) in PgVV-16T and PgVV-14T. The sequences of the TIRs were subdivided into (near) identical units based on their appearance in the four flanking regions. Note that the ends of the PgVV-14T genome could not be fully assembled and are thus truncated. Due to the lack of the unit F in 5’ TIR of PgVV-14T the region of the TIRs in this isolate (shown with a green frame) is shorter than in PgVV-16T. Locations of the first and the last ORFs are shown with red frames. Positions of the dinucleotide variation in units of type A are indicated.

**Extended Data Figure 2.**
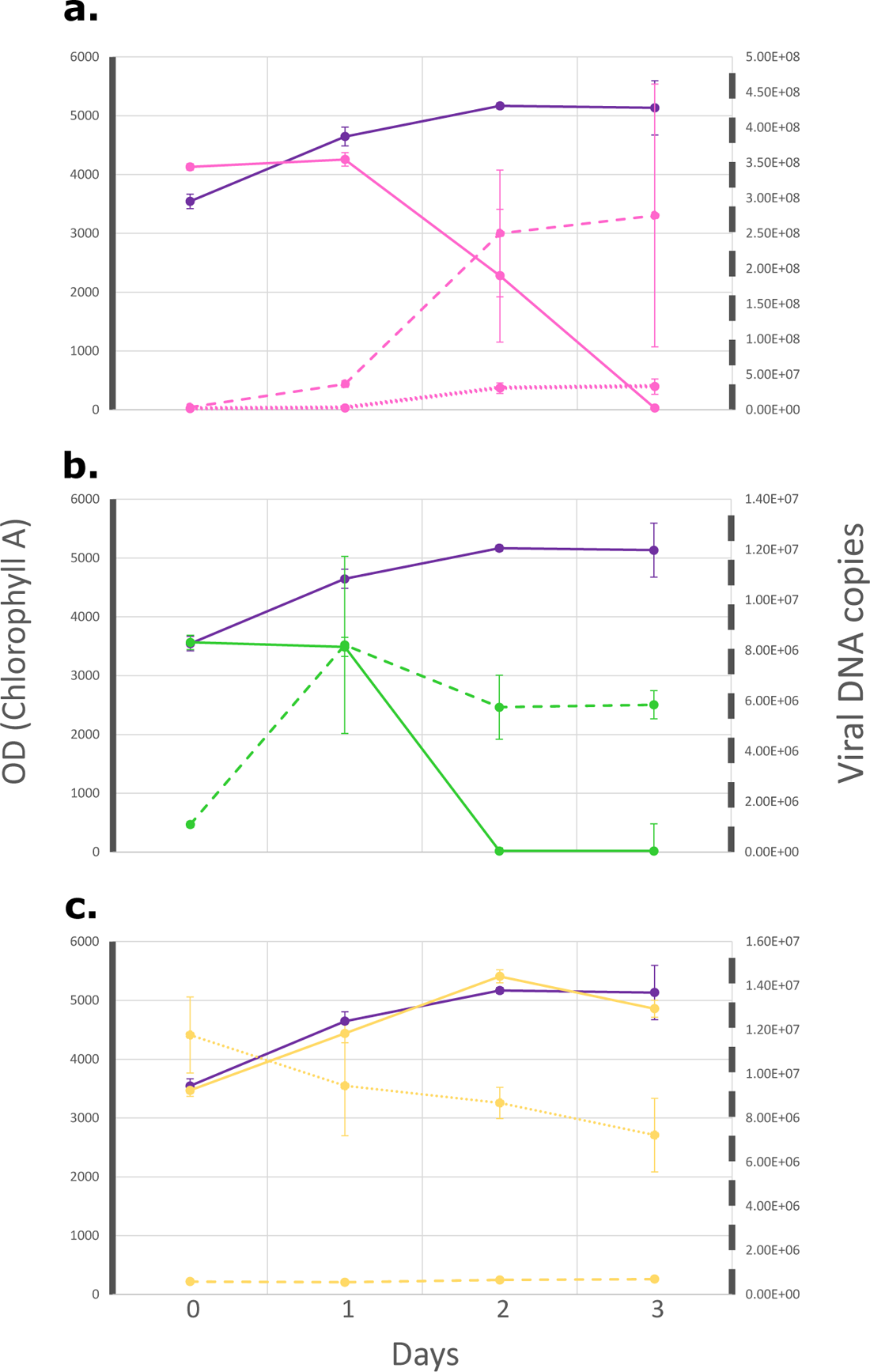
PgVV is a virophage. Infection experiments on *P. globosa* cultures infected with **a.** a mixed PgV/PgVV lysate, **b.** a pure PgV lysate, **c.** PgVV only. Purple lines denote the uninfected, control culture; full lines the cell survival measured by OD of chlorophyll A. Viral abundances were calculated by qPCR and are marked as dashed lines for PgV and dotted lines for PgVV.

**Extended Data Figure 3.**
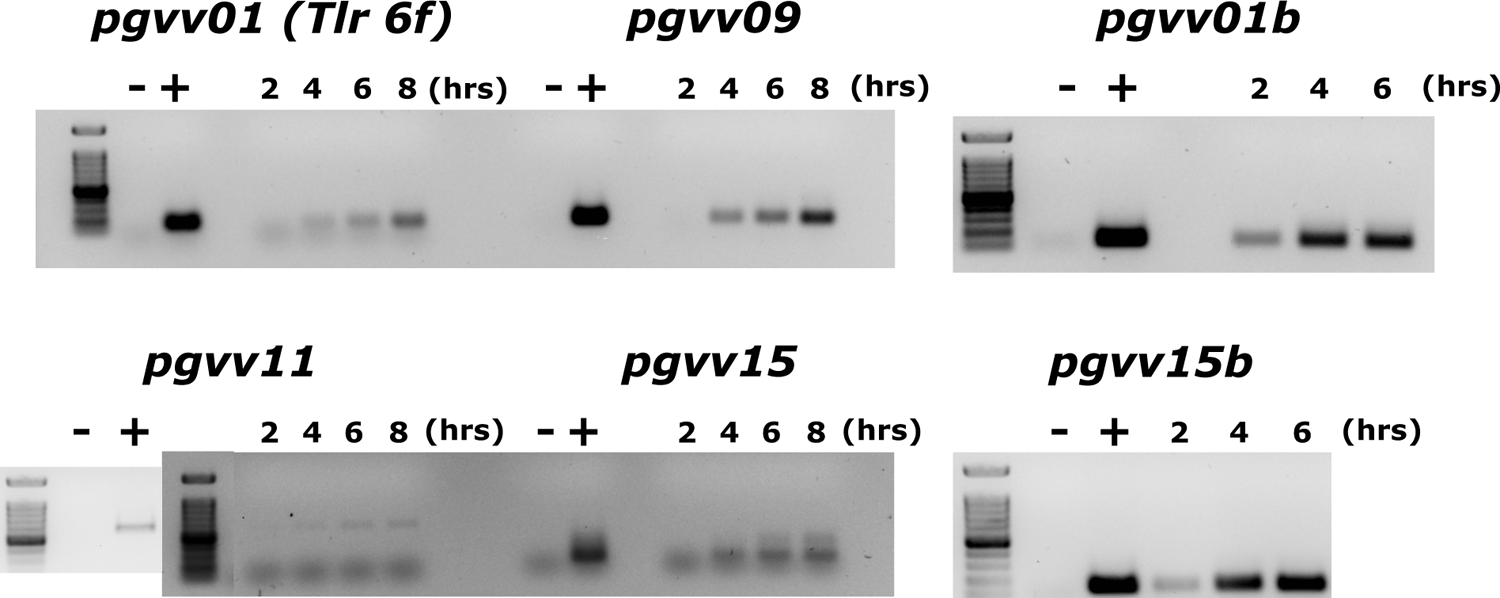
PgVV ORFs not detected by proteomics are transcribed during infection. PCR on cDNA for the six ORFs with no significant hits in the proteomics analyses. -, non-template control; +, positive control (PgVV DNA); samples were collected 2,4 and 6 hrs post-infection.

**Extended Data Figure 4.**
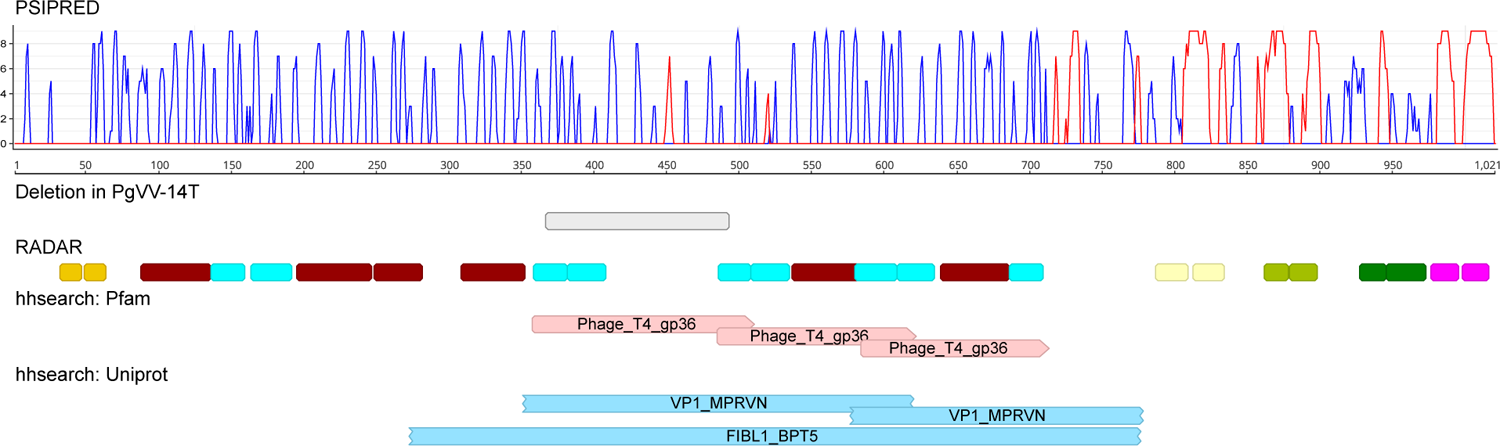
Secondary structure and domain composition of the protein coded by PgVV-16T ORF PGVV_00014. *First track:* per-position PSIPRED secondary structure prediction with blue lines corresponding to beta sheets and red lines to alpha helices (coils not shown), Y axis reflects confidence values (0-9). *Second track:* position of the large deletion in PgVV-14T. *Third track:* amino acid repeats discovered with RADAR with each color corresponding to a repeat type. *Fourth track:* locations of the hhsearch matches to Pfam profile PF03903 (bacteriophage T4 tail-fiber protein gp36) when searched against the Pfam database distributed with HH-Suite. *Fifth track:* hhsearch matches to Uniprot records: VP1_MPRVN – protein VP1 of Micromonas pusilla reovirus (Q1I0V1); FIBL1_BPT5 – L-shaped tail-fiber protein pb1 of Escherichia phage T5 (P13390).

**Extended Data Figure 5.**
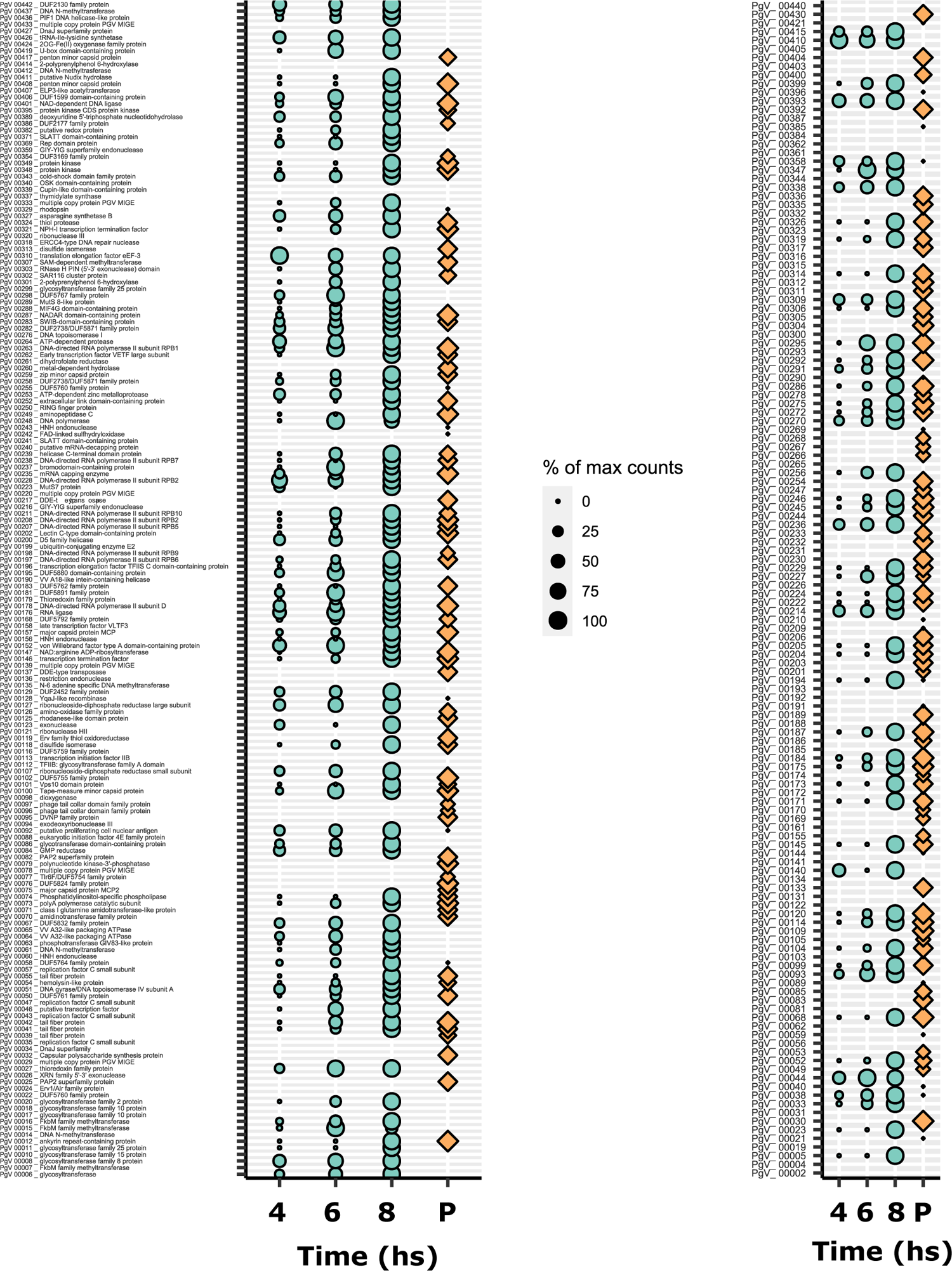
Proteomic analysis of PgV particles and during infection. Proteins found by mass spectrometry in purified PgV viral particles (rhomboids) and 4, 6 and 8 hours post-infection (circles). P - Purified viral particles. Relative quantification as described in the methods section. Dots mark samples where relevant peptides were found, but below the significance threshold. Right column includes all detected uncharacterized proteins.

**Extended Data Figure 6.**
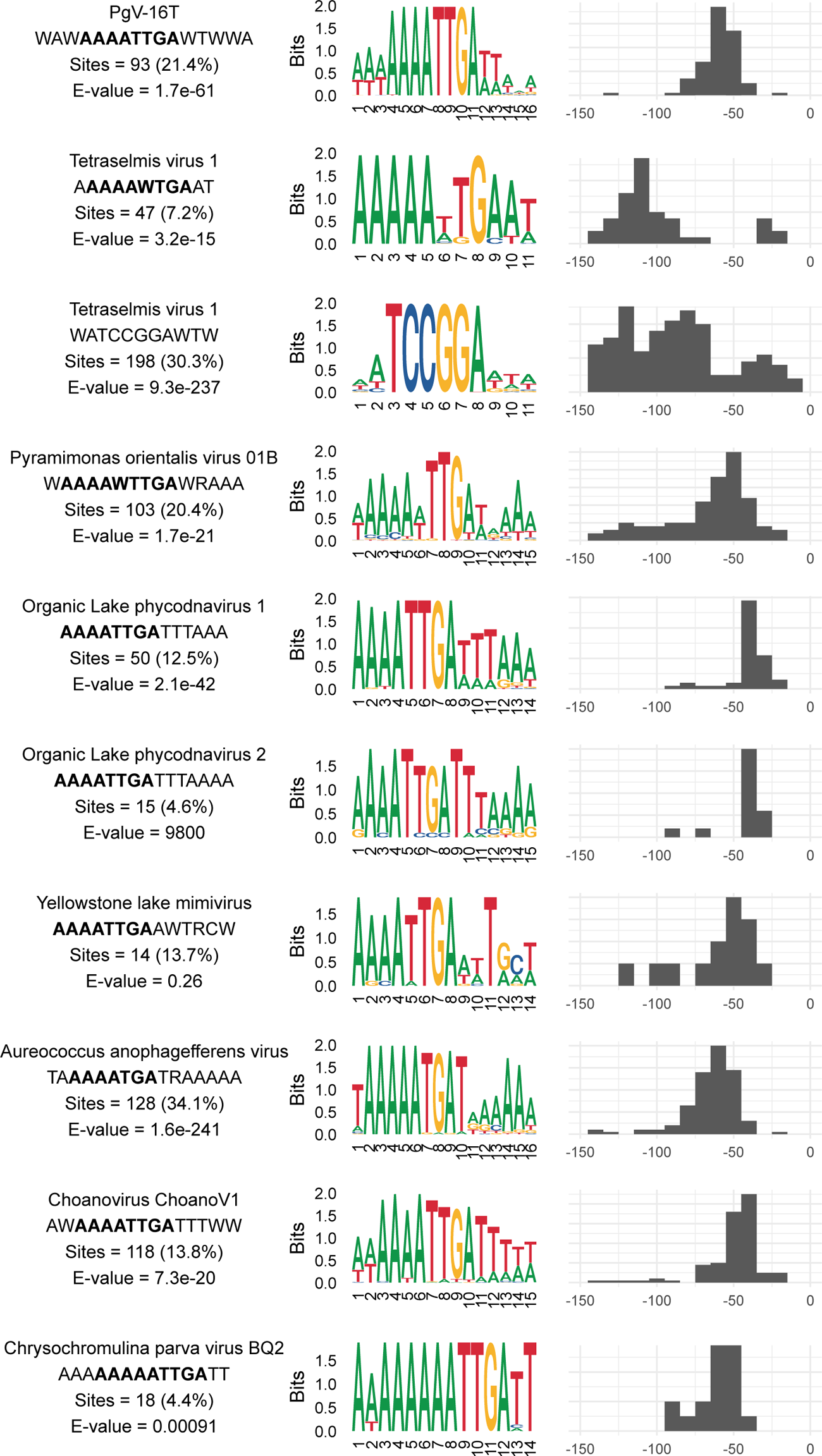
Early mimiviral promoter motif among mesomimiviruses. Results of the MEME search for common motifs in sequences upstream of ORFs in mesomimiviruses. Only motifs fitting the pattern WWWWWTGW are shown, supplemented by the unusually high-frequency palindromic motif TCCGGA of *Tetraselmis* virus 1. For each motif, a consensus sequence, number of sites and E-value are provided (to the left). Per-position weblogos and frequency distribution of distances from the start codon are shown in the middle and to the right.

**Extended Data Figure 7.**
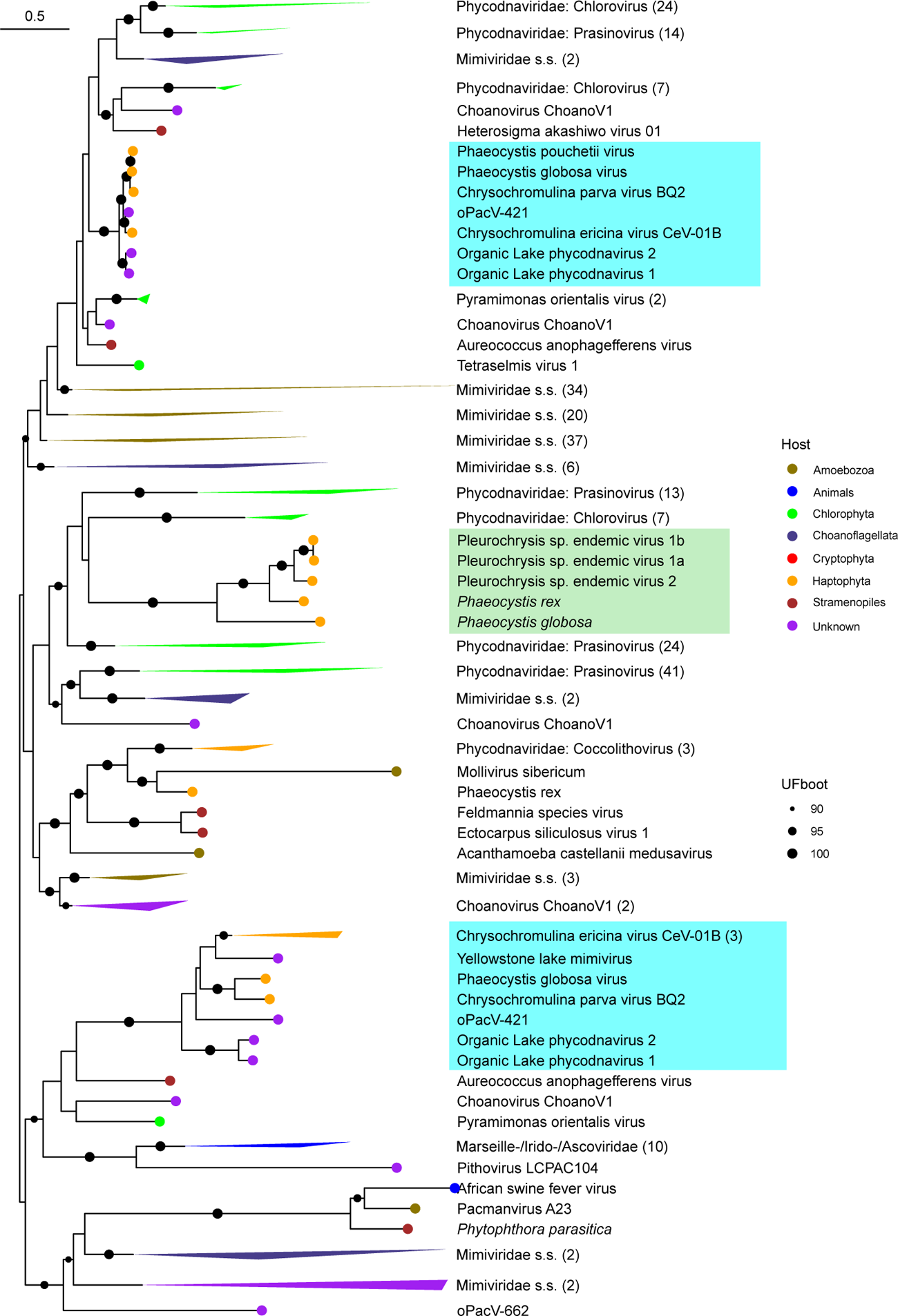
Phylogenetic analysis of MCPs from NCLDVs and NCLDV-like dwarf viruses. The clade including the MCPs of NCLDV-like dwarf viruses (NDDVs) is highlighted in green and MCPs appearing in mesomimiviral genes are highlighted in cyan. The tree is midpoint-rooted. Host groups are indicated when known. Numbers of sequences for collapsed clades are shown in parentheses.

**Extended Data Figure 8.**
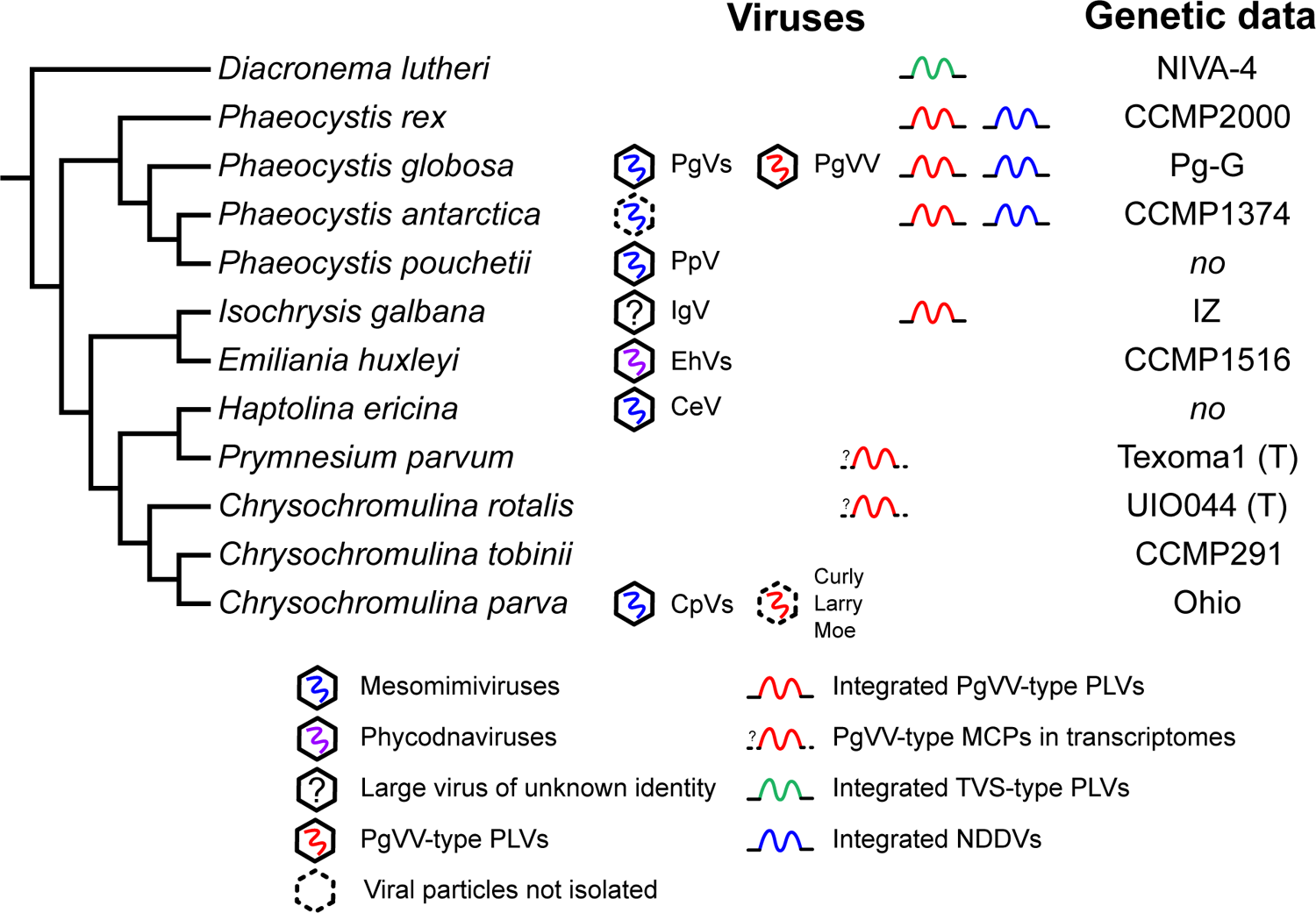
Distribution of NCLDVs, PLVs and endemic viruses among haptophytes. The cladogram is after ^84–86^. Strains available with genomic data suitable for analysis of integrated viruses are indicated (asterisks mark strains for which only transcriptomes are available). CeV – Chrysochromulina ericina virus, CpVs – *Chrysochromulina parva* viruses; EhVs – *Emiliania huxleyi* viruses, IgV – Isochrysis galbana virus ^87^, “PaV” – “Phycodnaviridae Antarctica virus” (mesomimivirus hypothesized to infect *P. antarctica* ^88^), PgVs – *Phaeocystis globosa* viruses, PpV – Phaeocystis pouchetii virus.

**Extended Data Figure 9.**
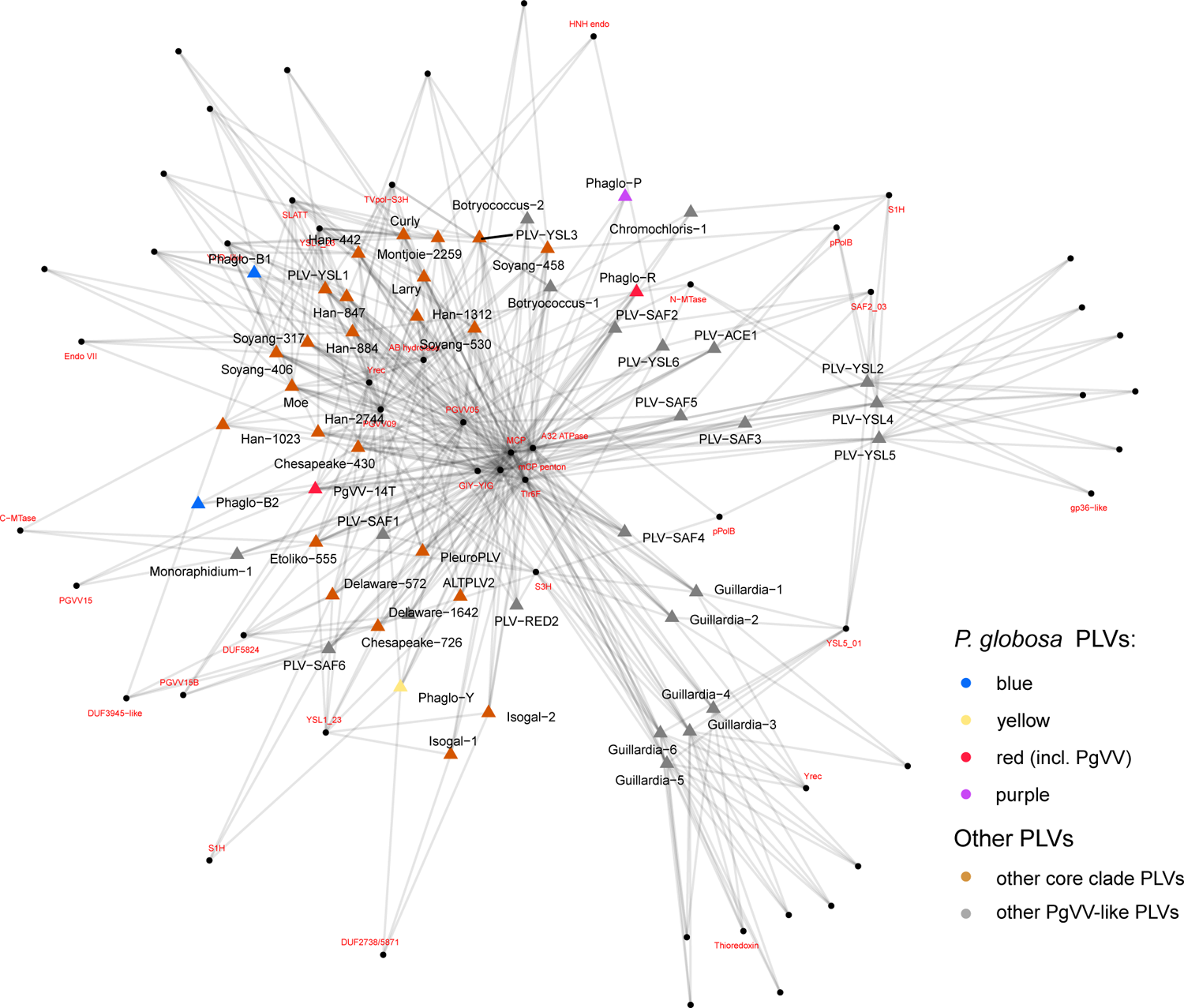
Bipartite network of gene cluster sharing between PgVV-group PLVs. Triangles represent individual PLV genomes. Genes were clustered based on profile-profile matches (see Materials and Methods) and each cluster is represented as a Red labels are provided for clusters that could be associated with widespread and/or functional families (see Table S1 for definitions of the widespread families).

**Extended Data Figure 10.**
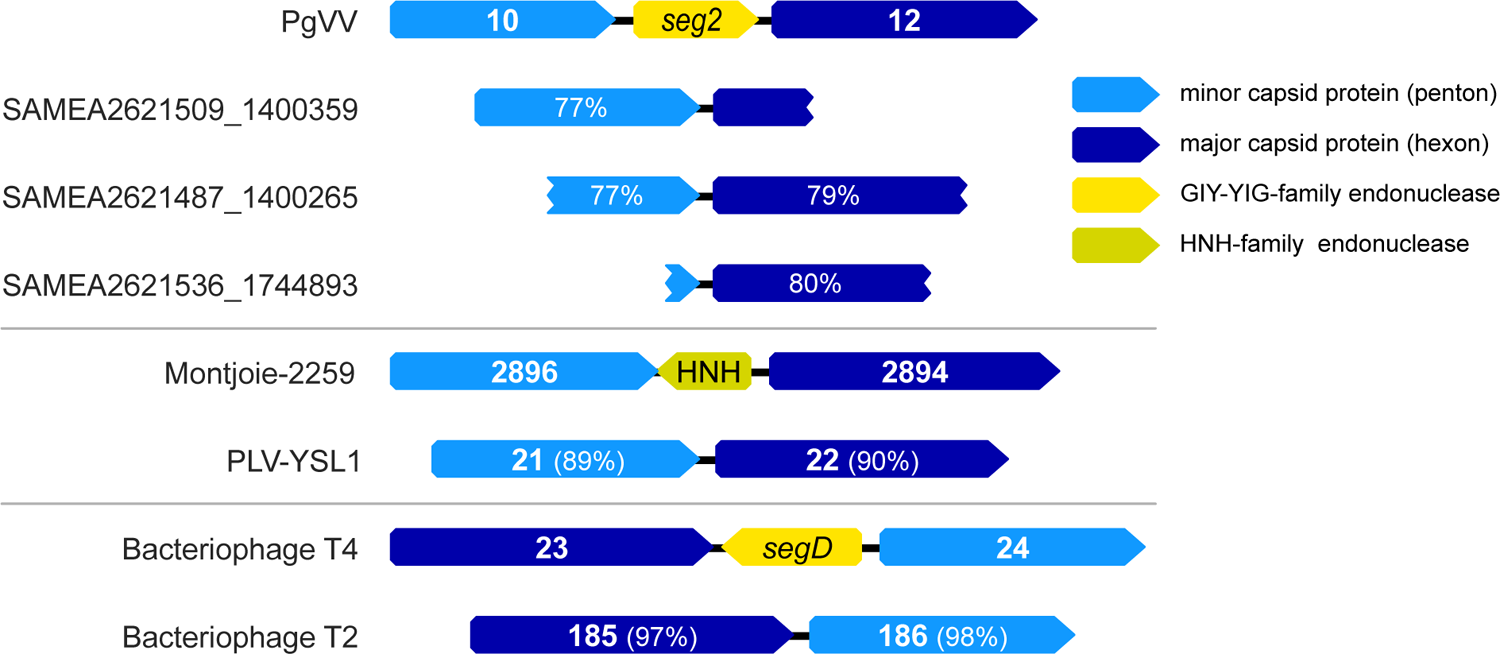
Incidence and mobility of genes coding for putative non-intronic homing endonucleases located between genes for capsid proteins. From top to bottom: GIY-YIG endonuclease gene *seg2* between mCP and MCP genes (ORFs *pgvv10* and *pgvv12*) in PgVV and its absence in closely related viruses as evidenced by the three metagenomic contigs; HNH endonuclease gene in the PLV Montjoie2259 and its lack in members of the same subgroup as exemplified by PLV-YSL1; a similar case of *segD*, a gene for a GIY-YIG-family endonuclease located between genes coding for the hexon and penton proteins in *Escherichia virus T4* present in bacteriophage T4 but absent from bacteriophage T2. ORF numbers are provided, percentages show similarity at the DNA level.

## References

1. Koonin, E. V. & Dolja, V. V. Virus World as an Evolutionary Network of Viruses and Capsidless Selfish Elements. Microbiol. Mol. Biol. Rev. MMBR 78, 278–303 (2014).

2. Pritham, E. J., Putliwala, T. & Feschotte, C. Mavericks, a novel class of giant transposable elements widespread in eukaryotes and related to DNA viruses. Gene 390, 3–17 (2007).

3. Kapitonov, V. V. & Jurka, J. Self-synthesizing DNA transposons in eukaryotes. Proc. Natl. Acad. Sci. 103, 4540–4545 (2006).

4. Krupovic, M. & Koonin, E. V. Polintons: a hotbed of eukaryotic virus, transposon and plasmid evolution. Nat. Rev. Microbiol. 13, 105–115 (2015).

5. Koonin, E. V., Krupovic, M. & Yutin, N. Evolution of double-stranded DNA viruses of eukaryotes: from bacteriophages to transposons to giant viruses. Ann. N. Y. Acad. Sci. 1341, 10–24 (2015).

6. Yutin, N., Raoult, D. & Koonin, E. V. Virophages, polintons, and transpovirons: a complex evolutionary network of diverse selfish genetic elements with different reproduction strategies. Virol. J. 10, 158 (2013).

7. Krupovic, M., Bamford, D. H. & Koonin, E. V. Conservation of major and minor jelly-roll capsid proteins in Polinton (Maverick) transposons suggests that they are bona fide viruses. Biol. Direct 9, 6 (2014).

8. Yutin, N., Shevchenko, S., Kapitonov, V., Krupovic, M. & Koonin, E. V. A novel group of diverse Polinton-like viruses discovered by metagenome analysis. BMC Biol. 13, 95 (2015).

9. Bellas, C. M. & Sommaruga, R. Polinton-like viruses are abundant in aquatic ecosystems. Microbiome 9, 13 (2021).

10. Pagarete, A., Grébert, T., Stepanova, O., Sandaa, R.-A. & Bratbak, G. Tsv-N1: A Novel DNA Algal Virus that Infects Tetraselmis striata. Viruses 7, 3937–3953 (2015).

11. Bekliz, M., Colson, P. & La Scola, B. The Expanding Family of Virophages. Viruses 8, 317 (2016).

12. Fischer, M. G. The Virophage Family Lavidaviridae. Curr. Issues Mol. Biol. 1–24 (2021) doi:10.21775/cimb.040.001.

13. Desnues, C. et al. Provirophages and transpovirons as the diverse mobilome of giant viruses. Proc. Natl. Acad. Sci. 109, 18078–18083 (2012).

14. Campos, R. K. et al. Samba virus: a novel mimivirus from a giant rain forest, the Brazilian Amazon. Virol. J. 11, 95 (2014).

15. Gaia, M. et al. Broad spectrum of mimiviridae virophage allows its isolation using a mimivirus reporter. PLoS One 8, e61912 (2013).

16. Hackl, T., Duponchel, S., Barenhoff, K., Weinmann, A. & Fischer, M. G. Virophages and retrotransposons colonize the genomes of a heterotrophic flagellate. Elife 10, e72674 (2021).

17. Yau, S. et al. Virophage control of antarctic algal host-virus dynamics. Proc Natl Acad Sci 108, 6163–8 (2011).

18. Gong, C. et al. Novel Virophages Discovered in a Freshwater Lake in China. Front. Microbiol. 7, 5 (2016).

19. Zhou, J. et al. Three Novel Virophage Genomes Discovered from Yellowstone Lake Metagenomes. J. Virol. 89, 1278–1285 (2014).

20. Yutin, N., Kapitonov, V. V. & Koonin, E. V. A new family of hybrid virophages from an animal gut metagenome. Biol. Direct 10, 19 (2015).

21. Stough, J. M. A. et al. Genome and Environmental Activity of a Chrysochromulina parva Virus and Its Virophages. Front. Microbiol. 10, 703 (2019).

22. La Scola, B. et al. The virophage as a unique parasite of the giant Mimivirus. Nature 455, 100–4 (2008).

23. Fischer, M. G. & Suttle, C. A. A Virophage at the Origin of Large DNA Transposons. Science 332, 231–234 (2011).

24. Gaia, M. et al. Zamilon, a Novel Virophage with Mimiviridae Host Specificity. PLoS One 9, e94923 (2014).

25. Mougari, S. et al. Guarani Virophage, a New Sputnik-Like Isolate From a Brazilian Lake. Front. Microbiol. 10, (2019).

26. Sheng, Y., Wu, Z., Xu, S. & Wang, Y. Isolation and Identification of a Large Green Alga Virus (Chlorella Virus XW01) of Mimiviridae and Its Virophage (Chlorella Virus Virophage SW01) by Using Unicellular Green Algal Cultures. J. Virol. 96, e02114–21 (2022).

27. Santini, S. et al. Genome of Phaeocystis globosa virus PgV-16T highlights the common ancestry of the largest known DNA viruses infecting eukaryotes. Proc Natl Acad Sci 110, 10800–10805 (2013).

28. Brussaard, C. P. D., Kuipers, B. & Veldhuis, M. J. W. A mesocosm study of Phaeocystis globosa population dynamics - 1. Regulatory role of viruses in bloom. Harmful Algae 4, 859–874 (2005).

29. Baudoux, A. C., Noordeloos, A. A. M., Veldhuis, M. J. W. & Brussaard, C. P. D. Virally induced mortality of Phaeocystis globosa during two spring blooms in temperate coastal waters. Aquat. Microb. Ecol. 44, 207–217 (2006).

30. Baudoux, A. C. & Brussaard, C. P. D. Characterization of different viruses infecting the marine harmful algal bloom species Phaeocystis globosa. Virology 341, 80–90 (2005).

31. Tarutani, K., Nagasaki, K. & Yamaguchi, M. Virus adsorption process determines virus susceptibility in Heterosigma akashiwo (Raphidophyceae). Aquat. Microb. Ecol. 42, 209–213 (2006).

32. Gann, E. R., Gainer, P. J., Reynolds, T. B. & Wilhelm, S. W. Influence of light on the infection of Aureococcus anophagefferens CCMP 1984 by a “giant virus”. PLoS ONE 15, (2020).

33. Van Etten, J. L., Burbank, D. E., Xia, Y. & Meints, R. H. Growth cycle of a virus, PBCV-1, that infects Chlorella-like algae. Virology 126, 117–125 (1983).

34. Boyer, M. et al. Mimivirus shows dramatic genome reduction after intraamoebal culture. Proc. Natl. Acad. Sci. 108, 10296–10301 (2011).

35. Desnues, C. & Raoult, D. Inside the Lifestyle of the Virophage. Intervirology 53, 293–303 (2010).

36. Sobhy, H., Scola, B. L., Pagnier, I., Raoult, D. & Colson, P. Identification of giant Mimivirus protein functions using RNA interference. Front. Microbiol. 6, (2015).

37. Fischer, M. G. & Hackl, T. Host genome integration and giant virus-induced reactivation of the virophage mavirus. Nature 540, 288–291 (2016).

38. Wodarz, D. Evolutionary dynamics of giant viruses and their virophages. Ecol. Evol. 3, 2103–2115 (2013).

39. Farr, G. A., Zhang, L. & Tattersall, P. Parvoviral virions deploy a capsid-tethered lipolytic enzyme to breach the endosomal membrane during cell entry. Proc. Natl. Acad. Sci. 102, 17148–17153 (2005).

40. Suhre, K., Audic, S. & Claverie, J.-M. Mimivirus gene promoters exhibit an unprecedented conservation among all eukaryotes. Proc Natl Acad Sci 102, 14689–14693 (2005).

41. Legendre, M. et al. mRNA deep sequencing reveals 75 new genes and a complex transcriptional landscape in Mimivirus. Genome Res. 20, 664–674 (2010).

42. Smith, D. R., Arrigo, K. R., Alderkamp, A.-C. & Allen, A. E. Massive difference in synonymous substitution rates among mitochondrial, plastid, and nuclear genes of Phaeocystis algae. Mol. Phylogenet. Evol. 71, 36–40 (2014).

43. Krupovic, M., Kuhn, J. H. & Fischer, M. G. A classification system for virophages and satellite viruses. Arch. Virol. 161, 233–247 (2016).

44. Suplatov, D. A., Besenmatter, W., Svedas, V. K. & Svendsen, A. Bioinformatic analysis of alpha/beta-hydrolase fold enzymes reveals subfamily-specific positions responsible for discrimination of amidase and lipase activities. Protein Eng. Des. Sel. 25, 689–697 (2012).

45. Burt, A. & Koufopanou, V. Homing endonuclease genes: the rise and fall and rise again of a selfish element. Curr. Opin. Genet. Dev. 14, 609–615 (2004).

46. Chase, E., Desnues, C. & Blanc, G. Integrated viral elements unveil the dual lifestyle of Tetraselmis spp. polinton-like viruses. http://biorxiv.org/lookup/doi/10.1101/2022.05.02.489867(2022) doi:10.1101/2022.05.02.489867.

47. Campbell, S., Aswad, A. & Katzourakis, A. Disentangling the origins of virophages and polintons. Curr. Opin. Virol. 25, 59–65 (2017).

48. Koonin, E. V., Senkevich, T. G. & Dolja, V. V. The ancient Virus World and evolution of cells. Biol. Direct 1, 29 (2006).

49. Guillard, R. R. L. Culture of Marine Invertebrate Animals: Proceedings — 1st Conference on Culture of Marine Invertebrate Animals Greenport. (Springer US, 1975). doi:10.1007/978-1-4615-8714-9.

50. Cottrell, M. & Suttle, C. Wide-spread occurrence and clonal variation in viruses which cause lysis of a cosmopolitan, eukaryotic marine phytoplankter Micromonas pusilla. Mar. Ecol. Prog. Ser. 78, 1–9 (1991).

51. Sullivan, M. B. DNA Extraction of Cesium Chloride-Purified Viruses using Wizard Prep Columns. (2016). doi: dx.doi.org/10.17504/protocols.io.c26yhd

52. González-Domínguez, J. & Schmidt, B. ParDRe: faster parallel duplicated reads removal tool for sequencing studies. Bioinformatics 32, 1562–1564 (2016).

53. Krueger, F., James, F., Ewels, P., Afyounian, E. & Schuster-Boeckler, B. FelixKrueger/TrimGalore: v0.6.7 - DOI via Zenodo. (2021) doi:10.5281/zenodo.5127899.

54. Bankevich, A. et al. SPAdes: A New Genome Assembly Algorithm and Its Applications to Single-Cell Sequencing. J. Comput. Biol. 19, 455–477 (2012).

55. Deng, Z. & Delwart, E. ContigExtender: a new approach to improving de novo sequence assembly for viral metagenomics data. BMC Bioinformatics 22, 119 (2021).

56. Schneider, C. A., Rasband, W. S. & Eliceiri, K. W. NIH Image to ImageJ: 25 years of image analysis. Nat. Methods 9, 671–675 (2012).

57. Patel, A. et al. Virus and prokaryote enumeration from planktonic aquatic environments by epifluorescence microscopy with SYBR Green I. Nat. Protoc. 2, 269–276 (2007).

58. Bolte, S. & Cordelières, F. P. A guided tour into subcellular colocalization analysis in light microscopy. J. Microsc. 224, 213–232 (2006).

59. Brussaard, C. P. D. Optimization of Procedures for Counting Viruses by Flow Cytometry. Appl. Environ. Microbiol. 70, 1506–1513 (2004).

60. Kirzner, S., Barak, E. & Lindell, D. Variability in progeny production and virulence of cyanophages determined at the single-cell level. Environ. Microbiol. Rep. 8, 605–613 (2016).

61. Ziv, I. et al. A Perturbed Ubiquitin Landscape Distinguishes Between Ubiquitin in Trafficking and in Proteolysis. Mol. Cell. Proteomics MCP 10, M111.009753 (2011).

62. HaileMariam, M. et al. S-Trap, an Ultrafast Sample-Preparation Approach for Shotgun Proteomics. J. Proteome Res. 17, 2917–2924 (2018).

63. Rappsilber, J., Mann, M. & Ishihama, Y. Protocol for micro-purification, enrichment, pre-fractionation and storage of peptides for proteomics using StageTips. Nat. Protoc. 2, 1896–1906 (2007).

64. Kong, A. T., Leprevost, F. V., Avtonomov, D. M., Mellacheruvu, D. & Nesvizhskii, A. I. MSFragger: ultrafast and comprehensive peptide identification in mass spectrometry–based proteomics. Nat. Methods 14, 513–520 (2017).

65. Li, W. & Godzik, A. Cd-hit: a fast program for clustering and comparing large sets of protein or nucleotide sequences. Bioinformatics 22, 1658–1659 (2006).

66. Lechner, M. et al. Proteinortho: Detection of (Co-)orthologs in large-scale analysis. BMC Bioinformatics 12, 124 (2011).

67. Buchfink, B., Reuter, K. & Drost, H.-G. Sensitive protein alignments at tree-of-life scale using DIAMOND. Nat. Methods 18, 366–368 (2021).

68. Mirdita, M. et al. ColabFold - Making protein folding accessible to all. 2021.08.15.456425 Preprint at https://doi.org/10.1101/2021.08.15.456425 (2022).

69. Jumper, J. et al. Highly accurate protein structure prediction with AlphaFold. Nature 596, 583–589 (2021).

70. O’Connell, J. et al. NxTrim: optimized trimming of Illumina mate pair reads. Bioinformatics 31, 2035–2037 (2015).

71. Li, D., Liu, C.-M., Luo, R., Sadakane, K. & Lam, T.-W. MEGAHIT: an ultra-fast single-node solution for large and complex metagenomics assembly via succinct de Bruijn graph. Bioinformatics 31, 1674–1676 (2015).

72. Luo, R. et al. SOAPdenovo2: an empirically improved memory-efficient short-read de novo assembler. GigaScience 1, 2047-217X-1–18 (2012).

73. Chevreux, B., Wetter, T. & Suhai, S. Genome Sequence Assembly Using Trace Signals and Additional Sequence Information. in Proceedings of the German Conference on Bioinformatics, GCB 1999, October 4-6, 1999, Hannover, Germany 45–56 (1999).

74. Walker, B. J. et al. Pilon: An Integrated Tool for Comprehensive Microbial Variant Detection and Genome Assembly Improvement. PLoS One 9, e112963 (2014).

75. Eddy, S. R. Accelerated Profile HMM Searches. PLoS Comput. Biol. 7, e1002195 (2011).

76. Minh, B. Q. et al. IQ-TREE 2: New Models and Efficient Methods for Phylogenetic Inference in the Genomic Era. Mol. Biol. Evol. 37, 1530–1534 (2020).

77. Hoang, D. T., Chernomor, O., von Haeseler, A., Minh, B. Q. & Vinh, L. S. UFBoot2: Improving the Ultrafast Bootstrap Approximation. Mol. Biol. Evol. 35, 518–522 (2018).

78. Kozlov, A. M., Darriba, D., Flouri, T., Morel, B. & Stamatakis, A. RAxML-NG: a fast, scalable and user-friendly tool for maximum likelihood phylogenetic inference. Bioinformatics 35, 4453–4455 (2019).

79. Steinegger, M. & Söding, J. MMseqs2 enables sensitive protein sequence searching for the analysis of massive data sets. Nat. Biotechnol. 35, 1026–1028 (2017).

80. Steinegger, M. et al. HH-suite3 for fast remote homology detection and deep protein annotation. BMC Bioinformatics 20, 473 (2019).

81. Enright, A. J., Van Dongen, S. & Ouzounis, C. A. An efficient algorithm for large-scale detection of protein families. Nucleic Acids Res. 30, 1575–1584 (2002).

82. Zimmermann, L. et al. A Completely Reimplemented MPI Bioinformatics Toolkit with a New HHpred Server at its Core. J. Mol. Biol. 430, 2237–2243 (2018).

83. Heger, A. & Holm, L. Rapid automatic detection and alignment of repeats in protein sequences. Proteins Struct. Funct. Bioinforma. 41, 224–237 (2000).

84. Egge, E. S., Eikrem, W. & Edvardsen, B. Deep-branching Novel Lineages and High Diversity of Haptophytes in the Skagerrak (Norway) Uncovered by 454 Pyrosequencing. J. Eukaryot. Microbiol. 62, 121–140 (2015).

85. Hovde, B. T. et al. *Chrysochromulina*: Genomic assessment and taxonomic diagnosis of the type species for an oleaginous algal clade. Algal Res. 37, 307–319 (2019).

86. Andersen, R. A., Bailey, J. C., Decelle, J. & Probert, I. *Phaeocystis rex* sp. nov. (Phaeocystales, Prymnesiophyceae): a new solitary species that produces a multilayered scale cell covering. Eur. J. Phycol. 50, 207–222 (2015).

87. Stepanova, O. A. Black Sea algal viruses. *Russ*. J. Mar. Biol. 42, 123–127 (2016).

88. Alarcón-Schumacher, T., Guajardo-Leiva, S., Antón, J. & Díez, B. Elucidating Viral Communities During a Phytoplankton Bloom on the West Antarctic Peninsula. Front. Microbiol. 10, 1014 (2019).

