## Supplementary File 1 for "Infection cycle and phylogeny of the Polinton-like virus Phaeocystis globosa virus virophage-14T"

### This file includes:

Supplementary results

Table S1

References

### Other supplementary material:

**Supplementary File 2:** Primers list, *P. globosa*-PgV-PgVV infection dynamics,

**Supplementary File 3:** Genome comparison of -16T and -14T viral strains, virions measurements.

**Supplementary File 4:** Proteomics data, PCRs for poly-cystronic transcripts.

**Supplementary File 5:** Metagenomic and Metatranscriptomics assemblies analyzed in this project, integrated and standalone viruses index and sequences, PCR for *P. globosa* PLVs. Excel format.

**Supplementary File 6:** Cluster affiliation and best hhsearch Pfam hits for protein sequences from mesomimiviruses, virophages, PLVs and NCLDV-like dwarf viruses. Excel format.

**Supplementary File 7:** Annotated viruses and viral fragments found in algal genomes. GenBank format.

### Supplementary Results and Discussion

**PgV-14T and PgVV-14T genome sequencing.** In the framework of our work we sequenced the genome of the isolate PgV-14T. Notwithstanding a small number of unresolved repetitive regions, we produced a near-complete assembly of the PgV-14T isolate amounting to 459,420 bp improving upon the unpublished draft genome of PgV-14T (Genbank accession HQ634144.1). 81.75% of the 585,724 trimmed unique read pairs could be mapped back to the PgV-14T genome. Along the PgV's complete genome, we assembled a separate scaffold of 17,480 bp corresponding to a genome closely related to PgVV (Genbank accession KC662250.1). Complex interplay of different repeat types in the flanking TIR regions (see below) prevented us from assembling them completely, yet the scaffold extended further to a final length of 19,217 bp.

**Terminal repeat organization**. PgVV-14T and PgVV-16T differ significantly in in the TIR regions (Extended Data Fig. 1) with the same repeat unit types of 10-318 nt in length ordered differently in the two genomes: the order ABCADEF is characteristic to both TIRs of PgVV-16T contrasting with BCADCADEF found in the 3' TIR of PgVV-14T and […]ADCADE in the 5' TIR. In addition, the 3’ TIR of PgVV-14T contains an internal inverted repeat G of at least 77 nt overlapping with unit B which is absent from the assembly of PgVV-16T. Such a repeat has the propensity to build a hairpin structure, and the 5’ end of the 5’ TIR that could not be assembled likely has the same structure. The sequences of the repeat units are identical between the two PgVV isolates, with the exception of the 94-nt repeat unit A that contained the above-mentioned dinucleotide variation affecting ORF *pgvv16* (Extended Data Fig. 1, Supplementary File 3).

**Overview of the gene repertoire.** We annotated the PgVV genomes using a combination of de novo gene prediction tools, homology searches, as well as proteomic analysis and amplification of cDNA. The genome of PgVV-16T was reported to contain 16 ORFs located on the same strand[1] and given the high similarity with PgVV-14T the same ORF numeration could be adopted for both isolates, here referred to as *pgvv01-pgvv16*. Furthermore, a closer examination revealed the presence of two additional short ORFs in both genomes: *pgvv01b* and *pgvv15b*. Gene aliases were assigned to nine ORFs whose gene products were associated with predictable functions: *Seg1* (*pgvv02*, endonuclease), *Yrec* (*pgvv03*, Tyr-recombinase), *TVpol* (*pgvv04*, polymerase-primase), *ABH* (*pgvv06*, alpha/beta hydrolase, putative lipase), *A32* (*pgvv07*, packaging ATPase), *mCP* (*pgvv10*, minor capsid protein), *Seg2* (*pgvv11*, endonuclease), *MCP* (*pgvv12*, major capsid protein) and *Ltf* (*pgvv14*, L-shaped tail fiber-like protein). In addition, five more ORFs represented genes widespread among other PgVV-like PLVs but coding for proteins of unknown function: *Tlr6F* (*pgvv01*), *pgvv05*, *pgvv09*, *pgvv15* and *pgvv15b*. The remaining four genes were restricted to PgVV or its closest relatives: *pgvv01b*, *pgvv08*, *pgvv13* and *pgvv16*.

**Genes coding for structural proteins.** Putative structural proteins include the previously recognized gene products MCP (hexon) and mCP (penton) that are predicted to accept a double jelly-roll and a single jelly-roll fold, respectively (Fig. 3b), to which the gene product Ltf (PGVV14) can be added (see below). Curiously, as noticed before for PgVV-16T[1], in contrast to many previously reported PLVs and members of the *Lavidaviridae*, the genes *MCP* and *mCP* in PgVV are separated by an extra ORF *pgvv11*. This ORF appears to code for a Seg-like GIY-YIG endonuclease that we designate here as Seg2. Inspection of this region among closely related viruses yielded several metagenomic fragments that did not have the *Seg2* gene thus indicating its mobility (Extended Data Fig. 10). This situation is reminiscent of the *segD* gene in bacteriophage T4 that is similarly located between the genes coding for the hexon and the penton proteins gp24 and gp23 and is absent from related phages[2]. Most of the Seg-family endonucleases of phage T4 function as free-standing (intronless) site-specific homing endonucleases[2–4], although experimental evidence for the activity of SegD itself is lacking[5]. Among PgVV-type PLVs we identified a similar case in the PLV Montjoie-2259, with a gene for HNH-family homing endonuclease inserted between ORFs coding for mCP and MCP proteins (Extended Data Fig. 10). Seg2 in PgVV is thus probably a homing endonuclease as well with its recognition site lying inside one of the neighboring genes (genes for capsid proteins in particular are expected to provide relative DNA conservation necessary for recognition). The PgVV genomes also contain a second related *seg*-like gene, *Seg1*, bringing the total number of endonucleases to two out of the 18 genes.

The third gene for which a structural function can be postulated is *Ltf* (*pgvv14*), the longest gene in the PgVV genomes. Secondary structure and amino acid repeat analysis indicate that the bulk of the Ltf protein comprises repeats with unit size of 23-51 residues composed of short beta-sheets (Extended Data Fig. 4). Remote homology searches indicate that this region includes several overlapping domains of 130-153 residues with similarity to gp36 of T4 phage, a component of the long tail fibers[6]. Particular similarity to Ltf of PgVV is demonstrated by the trimeric L-shaped tail fiber protein pb1 of T5 phage that also contains multiple gp36-like domains and is responsible for binding to the surface of *E. coli* outer membrane[7]. The deletion observed in the *Ltf* gene in PgVV-14T corresponds to one entire gp36-like unit indicating that the mutation preserved the overall structure of the protein, yet the corresponding morphological structure might be shorter in PgVV-14T. Viruses with genes coding for similar proteins include other PgVV-type PLVs (Fig. 4, Extended Data Fig. 9) and unrelated viruses such as Micromonas pusilla reovirus and most interestingly - PgV. Proteins encoded by three PgV ORFs: *PGCG_00041*, *PGCG_00042* and *PGCG_00055* contain gp36-like repeats, although they are longer than Ltf and otherwise have variable domain composition. The PgV ORF *PGCG_00042* in particular contains repetitive DNA stretches with high similarity to *Ltf* (hence the definition “PGCG_00042-like protein” in [1]) indicative of a recent limited recombination between the two genes.

**PLVs in *Phaeocystis* genomes.** Five genomic fragments from *P. globosa* were found to contain at least three core genes of the PgVV-like core. However, since our assembly is rather incomplete, and based on the many MCP sequences we detected, we suspect there are many more PLVs from the different groups in the host genome. Besides assembly incompleteness, PLV degradation is in part responsible for the fragmented nature of the other PLVs as demonstrated by the presence of a reverse transcriptase gene at the 5’ flank of Phaglo-R that disrupted a helicase gene in this PLV (Fig. 4). Phaglo-Y seems to be a complete PLV, flanked with TIRs and including many homologous to core PgVV genes. PgVV-like PLVs were found in the genomes assemblies of all three *Phaeocystis* species for which genomic data are available. In our assembly of *P. globosa* the PLVs seem integrated to the genome. In total, four subclades of PgVV-like PLVs are detected in *Phaeocystis* genomes, each one with its own host range, which might mimic the variable host range of yet-uncultured giant viruses infecting *Phaeocystis* species. The purple (represented by Phaglo-P) and the red (Phaglo-R) groups are present only in *P. globosa,* the blue group (Phaglo-B1, -B2) is found in *P. globosa* and *P. antarctica*, while the yellow group (Phaglo-Y) is found also in *P. globosa*, *P. antarctica* and *P. rex*. The MCP phylogeny of those PLVs mirror the host species evolution[8], suggesting that the possible integration/coupling of the PLVs from the yellow group predated the divergence of *P. rex*, while the PLVs from the red and purple groups integrated after the separation of *P. globosa* and *P. antarctica*. Phaglo-G, whose MCP is of a distinct origin than the others seems to be directly related to giant viruses, and encode four out of five genes universally present among NCLDVs: MCP, ATPase, VLTF3 and SF3 helicase (in PsEV1b and PsEV2)[9], as well as other typical NCLDV genes, such as YqaJ recombinase and RuvC resolvase[9,10] (Fig. 4). No type B polymerase genes could be found, but Phaglo-G and PsEVunk instead contained a TVpol-S3H gene.

**Table S1. Conserved genes among PgVV-type PLVs**. For each gene a formal definition is given based on hhsearch matches against the HH-Suite Pfam database or on curated cluster members. Note that some genes encompass several clusters while others only only one. ORF numbers in the PgVV-16T and PgVV-14T genomes are provided for genes present in PgVV.

| Family | Definition | Predicted function | PgVV gene |
|---|---|---|---|
| MCP | Clusters matching PF09018 (Phage_Capsid_P3; PLVs) and PF04451 (Capsid_NCLDV; mesomimiviruses) or include proteins Mavirus MV18 and Sputnik V20 (virophages) | Major capsid protein (hexon) | *MCP* (*pgvv12*) |
| mCP penton | Clusters that include PGVV10 (PLVs), TvV-S1 TVSG_00014 and TVSG_00022 (PLVs), Mavirus MV17 (virophages), PgV PGCG_00408 and PGCG_00417 (mesomimiviruses) | Minor capsid protein penton | *mCP* (*pgvv10*) |
| A32 ATPase | The cluster with the best hit to PF04665 (Pox_A32) | Packaging ATPase | *A32* (*pgvv07*) |
| Tlr6F | Cluster matching PF19058 (DUF5754) | Unknown | *Tlr6F* (*pgvv01*) |
| PGVV05 | = *G. theta* protein C. Cluster of PGVV05 | Unknown | *pgvv05* |
| AB hydrolase | Clusters matching PF11187 (Mbeg1-like) and other alpha-beta hydrolase profiles | Includes lipases, but potentially also other α/β hydrolases | *ABH* (*pgvv06*) |
| PGVV09 | = Conservative 4 in [11]. Cluster of PGVV09 | Unknown | *pgvv09* |
| SLATT | Clusters matching PF18186 (SLATT_4). The biggest cluster includes *G. theta* protein G. Have two transmembrane helices. | Contain putative pore-forming domain | |
| Yrec | Clusters with matches to PF00589 (Phage_integrase) and related profiles | Tyr-recombinase/ integrase | *Yrec* (*pgvv03*) |
| YSL1_23 | = Conservative 3 in [11]. Cluster of YSL1_23 | Unknown | |
| GIY-YIG | Clusters that include standalone GIY-YIG-type endonucleases (matching PF19835 (SegE_GIY-YIG) and PF01541 (GIY-YIG)). | GIY-YIG-type endonucleases, some are homing endonucleases | *seg1* (*pgvv02*), *seg2* (*pgvv11*) |
| gp36-like | Clusters with domains having remote similarity to PF03903 (Phage_T4_gp36). Proteins have otherwise varying lengths and diverse domain composition. | Putative structural proteins | *pgvv14* |
| S3H | Multiple clusters matching SF3 helicase profiles, not fused with TVpol | Standalone superfamily 3 helicases | |

| | | | |
|---|---|---|---|
| pPolB | Clusters with best matches to PF03175 (DNA_pol_B_2) or fusion proteins containing domains with matches to PF02689 (Herpes_Helicase) and PF03175 (see [13]). | Protein-primed β-type DNA polymerase | |
| TVpol-S3H | Clusters that include proteins matching both PF00476 (DNA_pol_A) and PF04735 (Baculo_helicase). See [12]. | TVpol-type DNA polymerase | *TVpol* (*pgvv05*) |
| YSL5_01 | Cluster of YSL5_01 | Unknown | |
| S1H | Multiple clusters matching SF1 helicase profiles, not fused to pPolB domains | Standalone superfamily 1 helicases | |
| N-MTase | Clusters matching PF02384 (N6_Mtase), PF20473 (MmeI_Mtase), PF02086 (MethyltransfD12), PF13651 (EcoRI_methylase), PF05869 (Dam) | Putative DNA N-methyltransferases | |
| ac53-Zfp | Clusters matching PF05883 (Baculo_RING) | Contain RING-finger domains related to ubiquitin ligases | |
| DUF5824 | Clusters matching PF19141 (DUF5824) | Unknown | |
| SAF2_03 | Cluster of SAF2_03 | Unknown | |
| DUF2738/5871 | Clusters matching PF10927 (DUF2738) and PF19196 (DUF5871) | Unknown | |
| DUF3945-like | Cluster with weak similarity to PF13101 (DUF3945) | Unknown | |
| PGVV15B | Cluster of PGVV15B | Unknown | *pgvv15b* |
| TAGT | Clusters matching PF20267 (GREB1_C), members of the TET/JBP-associated glycosyltrasferase (TAGT) family | Putative DNA glycosyltransferase | |
| C-MTase | Clusters matching PF00145 (DNA_methylase) | Putative DNA C-methyltransferases | |

### References


1. Santini, S. *et al.* Genome of Phaeocystis globosa virus PgV-16T highlights the common ancestry of the largest known DNA viruses infecting eukaryotes. *Proc. Natl. Acad. Sci.* **110**, 10800–10805 (2013).
2. Edgell, D. R., Gibb, E. A. & Belfort, M. Mobile DNA elements in T4 and related phages. *Virol J* **7**, 290 (2010).
3. Edgell, D. R. Free-standing homing endonucleases of T-even phage: freeloaders or functionaries? in *Homing Endonucleases and Inteins* 147–160 (2005).
4. Belle, A., Landthaler, M. & Shub, D. A. Intronless homing: site-specific endonuclease SegF of bacteriophage T4 mediates localized marker exclusion analogous to homing endonucleases of group I introns. *Genes Dev.* **16**, 351–362 (2002).
5. Sokolov, A. S. *et al.* Phage T4 endonuclease SegD that is similar to group I intron endonucleases does not initiate homing of its own gene. *Virology* **515**, 215–222 (2018).
6. Islam, M. Z. *et al.* Molecular anatomy of the receptor binding module of a bacteriophage long tail fiber. *PLoS Pathogens* **15**, e1008193 (2019).
7. Zivanovic, Y. *et al.* Insights into Bacteriophage T5 Structure from Analysis of Its Morphogenesis Genes and Protein Components. *J. Virology* **88**, 1162–1174 (2014).
8. Medlin, L. & Zingone, A. A taxonomic review of the genus Phaeocystis. *Biogeochemistry* **83**, 3–18 (2007).
9. Yutin, N., Wolf, Y. I., Raoult, D. & Koonin, E. V. Eukaryotic large nucleo-cytoplasmic DNA viruses: Clusters of orthologous genes and reconstruction of viral genome evolution. *Virology Journal* **6**, 223 (2009).
10. Iyer, L. M., Balaji, S., Koonin, E. V. & Aravind, L. Evolutionary genomics of nucleo-cytoplasmic large DNA viruses. *Virus Research* **117**, 156–184 (2006).

11. Stough, J. M. A. *et al.* Genome and Environmental Activity of a Chrysochromulina parva Virus and Its Virophages. *Frontiers in Microbiology* **10**, 703 (2019).

12. Iyer, L. M., Abhiman, S. & Aravind, L. A new family of polymerases related to superfamily A DNA polymerases and T7-like DNA-dependent RNA polymerases. *Biol. Direct* **3**, 39 (2008).

13. Krupovic, M., Yutin, N. & Koonin, E. V. Fusion of a superfamily 1 helicase and an inactivated DNA polymerase is a signature of common evolutionary history of Polintons, polinton-like viruses, Tlr1 transposons and transpovirons. *Virus Evol* **2**, vew019 (2016).
